# Scalable transcription factor mapping uncovers the regulatory dynamics of natural and synthetic transcription factors in human T cell states

**DOI:** 10.1101/2025.10.09.681414

**Authors:** Riley D.Z. Mullins, Jesse Zaretsky, Emily Stoller, Michael Moore, Oliver Takacsi-Nagy, Oleg Shpynov, Remi Sampaleanu, Theodore L. Roth, Ansuman T. Satpathy, Robi D. Mitra, Sidharth V. Puram

## Abstract

Heterogeneous T cell states are critical in immune responses and have been explored by CRISPR-based and synthetic domain-swapped transcription factor (TF) screens, yielding novel insights and immunotherapeutics. However, a scalable strategy to map TFs in primary human T cells is lacking, which limits our understanding of the functions of critical TFs. We therefore adapted a transposon-based TF mapping strategy termed Calling Cards for primary human CD8 T cells, applying it to five key TFs with undefined binding sites in this cell type: TOX, TOX2, TCF7, SOX4, and RBPJ. To derive biological insights from these data, we developed an analytical framework to integrate TF binding with multi-omic sequencing data, revealing convergence of TOX and TCF7 binding at dynamic enhancers of memory CD8 T cells. We then identified TF co-bound gene programs related to memory and exhaustion states in addition to putative gene targets of known and unappreciated TF roles, including TOX binding at critical genes of both exhaustion and terminal effector memory differentiation. To further scale our TF analysis platform, we modified Calling Cards to create ***TFlex***: a method uniquely suited for multiplexed mapping of paralogous TFs. We applied TFlex to simultaneously map eight natural and domain-swapped TFs in primary human CD8 T cells, which demonstrated that domain-swapped TFs display emergent behavior in binding site selection and transcriptional effects on target genes that cannot be estimated as the sum of their constituent domains. Collectively, our data highlight the importance of scalable TF mapping in primary human T cells to elucidate TF function and the transcriptional regulation of cell states.

## INTRODUCTION

Cytotoxic (CD8+) T cells adopt heterogeneous cell states that are central to immune responses, including effector and memory phenotypes that help clear acute viral infections and exhaustion states that constrain chronic viral infections and cancer^1^. Although the application of CRISPR-based technologies in primary human T cell states has led to novel biological insights and clinically-approved therapies, the regulation of the broader transcriptional gene networks underlying these cell states remains poorly defined^2–6^. These networks are coordinated by transcription factors (TFs), which bind DNA to regulate gene expression^7^. Thus, it is critical to define TF binding sites in order to determine their gene targets and the larger gene programs they act upon to govern T cell states.

However, a scalable method to map TFs has yet to be applied in primary human T cells. Although high-throughput methods to map TFs have had success in other cell types, their use in primary human T cells has been limited by the absence of specific antibodies for crucial T cell TFs and challenges in delivering constructs into primary cells^8–10^. Hence, the binding sites and thus gene targets of many TFs important in regulating cell states are not defined in primary human T cells, including the quintessential example of TOX and TCF7 in exhaustion^11,12^. Furthermore, mapping synthetic domain-swapped TFs, which can enhance the anti-tumor functions of human T cells and may have therapeutic applications, is also a significant challenge with the current available methods^13^. Thus, there is a need for a new strategy to map natural and synthetic TFs at scale in primary human T cells to gain insight into TF-targeted gene programs and the mechanisms coordinating disease-relevant T cell states.

We therefore sought to develop a scalable platform to define TF binding sites and uncover their targeted gene programs in primary human T cells. We first adapted a transposon-based method to map TFs, termed Calling Cards (CCs)^14,15^, for use in primary human CD8 T cells. We applied this method to map five key TFs with undefined binding sites in this cell type: TOX, TOX2, TCF7, SOX4, and RBPJ. To derive biological insights from these data, we developed an analytical framework to integrate TF binding data with multi-omic sequencing data. With this approach, we found that TOX and TCF7 converge on enhancers of human memory CD8 T cells, despite their interplay classically thought to only be involved in exhaustion. We next used this framework on single cell RNA-sequencing (scRNA-seq) data to define TF co-bound gene programs that underlie core T cell states, including stemness, memory, exhaustion, and activation. While some programs had similar expression profiles, they were clearly distinguishable by differential TF binding, revealing the distinct TFs that regulate coinciding gene programs. These TF bound gene programs served to identify putative gene targets that mediate known TF functions, including the major roles of TCF7 in stemness and TOX in exhaustion, and unanticipated functions, such as involvement of TOX in terminal effector memory differentiation. Finally, to further scale our TF analysis platform, we modified the standard CCs method to create ***TFlex***: a method uniquely suited for multiplexed mapping of paralogous natural and synthetic TFs. To validate the multiplexed capacity of TFlex, we simultaneously mapped eight bZIP family TFs in primary human CD8 T cells, including JUN, FOSL1, BATF, and five Domain Engineered via Synthesis and Recombination (DESynR) TFs derived from recombined N-terminal, DNA-binding, and C-terminal domains of natural TFs. We found that these domain-swapped DESynR TFs can acquire emergent behavior in binding site selection and target gene effects that cannot be estimated as the sum of their constituent domains. Collectively, our data highlight the importance of scalable TF investigations in primary human T cells to elucidate how TFs regulate cell states and prioritize therapeutic strategies, particularly within synthetic biology.

## RESULTS

### Development of a Calling Cards assay for transposon-based mapping of transcription factors in primary human T cells

Our efforts to create a scalable platform for TF binding site analyses began with optimizing standard CCs in primary human T cells, which required overcoming the barrier of introducing vectors into these cells. The basis of CCs is the fusion of a TF to the hyperactive piggyBac transposase (HyPBase), which inserts a self-reporting transposon (SRT) into the genomic DNA at TF binding sites (**Fig. 1A**)^14^. The SRT is composed of a promoter and a marker gene flanked by terminal repeat sequences, which are recognized by HyPBase for transposition of the SRT into genomic DNA. The promoter produces mRNA of the marker gene and the junction of the SRT and genomic DNA, which is used to determine TF binding sites in sequencing data (**Fig 1A**). To use this assay in primary human T cells, we created an SRT encoding the elongation factor-1α (EF-1α) promoter and truncated CD34 (tCD34) as the marker gene, both of which are commonly used in human T cells (**Fig. 1A**)^16,17^. We sought to optimize an electroporation-based protocol for human CD8 T cells where cells were subjected to either *in vitro* acute activation for 72 hours or chronic activation for 11 days followed by electroporation of the CCs constructs (**Fig. 1B**). Cells were rested for 72 hours after electroporation to allow time for SRT insertions into genomic DNA (**Fig. 1B**). The chronic activation model was used to induce a state similar to exhaustion, and we confirmed it led to the expected decrease in inflammatory cytokine expression and increase in CD39 and TIM-3 expression (**Supplementary Fig. 1**). Because the efficient delivery of vectors in primary human T cells is challenging, we next performed extensive protocol development and optimization to deliver the HyPBase and SRT-tCD34 CCs constructs. We first tested plasmid purification methods and unexpectedly found that endotoxin-free preparations did not improve transposition efficiency in CD4 or CD8 primary human T cells (**Supplementary Fig. 2**). Nonetheless, we used endotoxin-free preparations as endotoxins may still activate innate immune pathways without affecting efficiency. We next tested alternative vector forms, including plasmid or minicircle SRT-tCD34 vectors and plasmid, minicircle, or mRNA HyPBase expression constructs (**Supplementary Fig. 3**). All told, co-electroporation of HyPBase mRNA with the minicircle SRT-tCD34 vector proved optimal, yielding an SRT insertion efficiency of approximately 90% in the acute and chronic activation conditions, relative to ∼50% for the pre-optimized approach using plasmid forms of each construct (**Fig. 1C, D**). Viability was only modestly reduced following electroporation (∼15% decrease in acute activation, ∼35% decrease in chronic activation) (**Fig. 1E**). Together, these data highlight our approach as an efficient means of delivering CCs vectors into primary human T cells, which is the foundation of scalable TF mapping by standard CCs and the derivative TFlex method.

**Figure 1.**
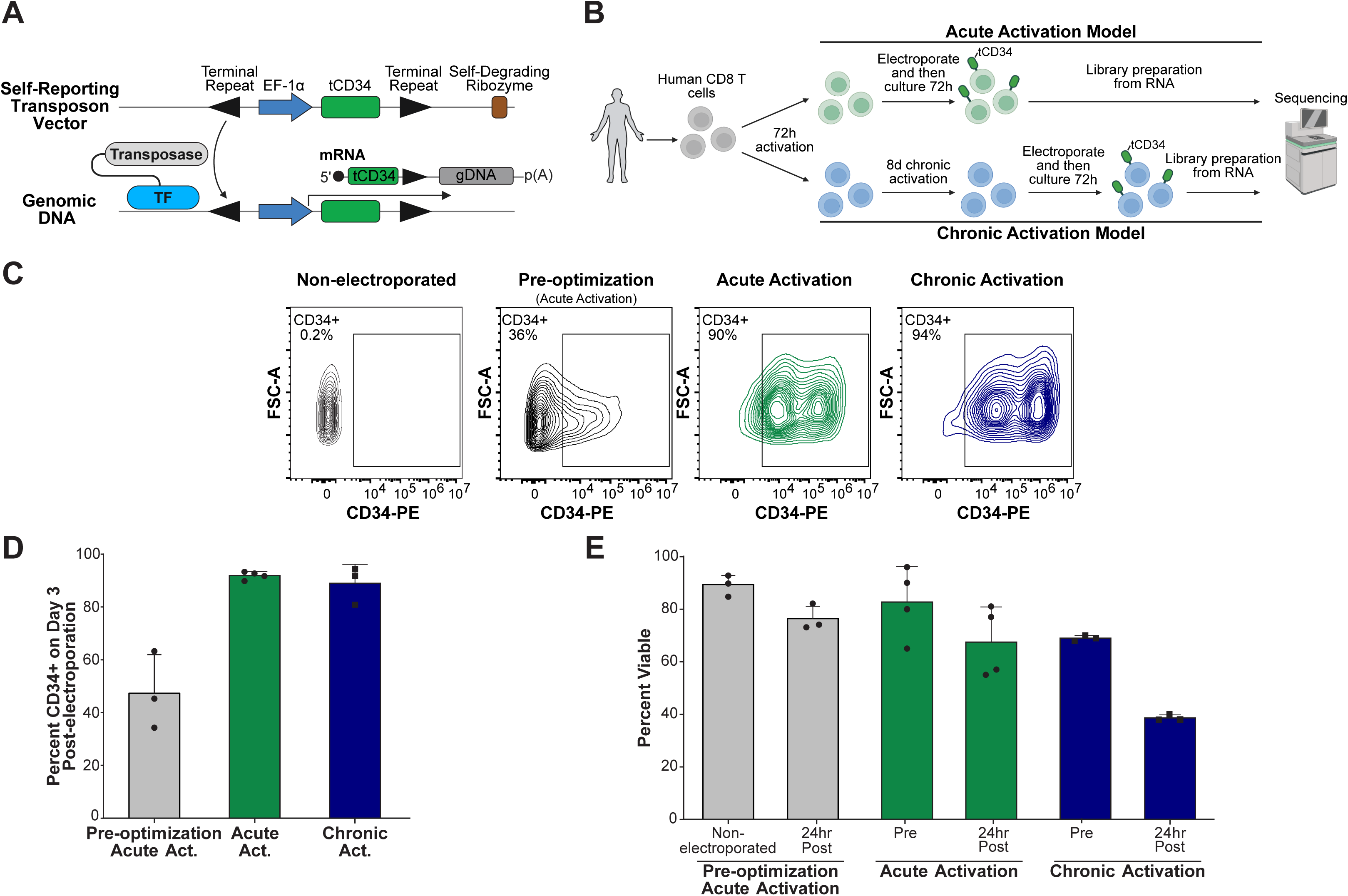
Development of a Calling Cards assay for primary human T cells. **(A)** Schematic of the CCs assay. The SRT is inserted into genomic DNA by a TF-transposase fusion protein at TF binding sites. The SRT consists of terminal repeat sequences flanking the EF-1α promoter and the marker gene, in this case truncated CD34 (tCD34). A self-degrading ribozyme sequence is downstream of the 3’ terminal repeat and serves to degrade transcripts arising from the SRT vector. When the SRT is integrated into the genomic DNA, transcripts are produced that include the junction of the terminal repeat and genomic DNA, which is detected by RNA-seq to define TF binding sites. **(B)** Schematic demonstrating the timing of *in vitro* primary human CD8 T cell activation, electroporation of CCs vectors, and RNA collection for CCs library preparation. **(C)** Representative contour plots of CD34 expression from the SRT-tCD34 transposon in CD8 T cells at 72 hours post-electroporation following *in vitro* acute or chronic activation using the final, optimized protocol or prior to full optimization (Pre-optimization). The pre-optimization data is from the experiment of Supplementary Figure 2. **(D)** Frequency of CD34+ CD8 T cells at 72 hours post-electroporation from the acute or chronic activation (Act.) model (n = 3 donors for pre-optimization; n = 4 donors otherwise, mean ± SD). The pre-optimization data is from the experiment of Supplementary Figure 2. **(E)** Frequency of viable cells pre-electroporation (Pre) and at 24 hours post-electroporation (24h Post) from the acute or chronic activation model (n = 3 donors for pre-optimization; n = 4 donors otherwise, mean ± SD). The pre-optimization data is from the experiment of Supplementary Figure 2.

### The Calling Cards assay robustly maps transcription factor binding sites in primary human CD8 T cells

With the delivery of the CCs vectors optimized, we next set out to leverage this approach to define putative target genes of previously uncharacterized TFs in primary human CD8 T cells. Specifically, we chose to map five TFs with critical roles in human CD8 T cells that, to our knowledge, have not been mapped in this cell type: TOX, TOX2, TCF7, SOX4, and RBPJ. We first confirmed that HyPBase in the TF-HyPBase fusions remained functional based on tCD34 expression from the SRT after electroporation (**Supplementary Fig. 4**). We then defined TF peaks, representing binding sites, as genomic regions with at least ten normalized SRT insertions and a log_2_ fold change in normalized SRT insertions greater than or equal to 1 relative to the unfused HyPBase background control (see **Methods**). As exemplified for the *IFNG* and *FAS* loci, these thresholds yielded peaks with a focused enrichment of SRT insertions that were demonstrably enriched above the background control, and most peaks were well above the log_2_ fold change threshold (**Fig. 2A, B**). In all, we defined a total of 6,500 to 15,500 peaks per TF, which represent their binding sites, in both acute and chronic activation conditions (**Fig. 2C; Supplementary Table 1**).

**Figure 2.**
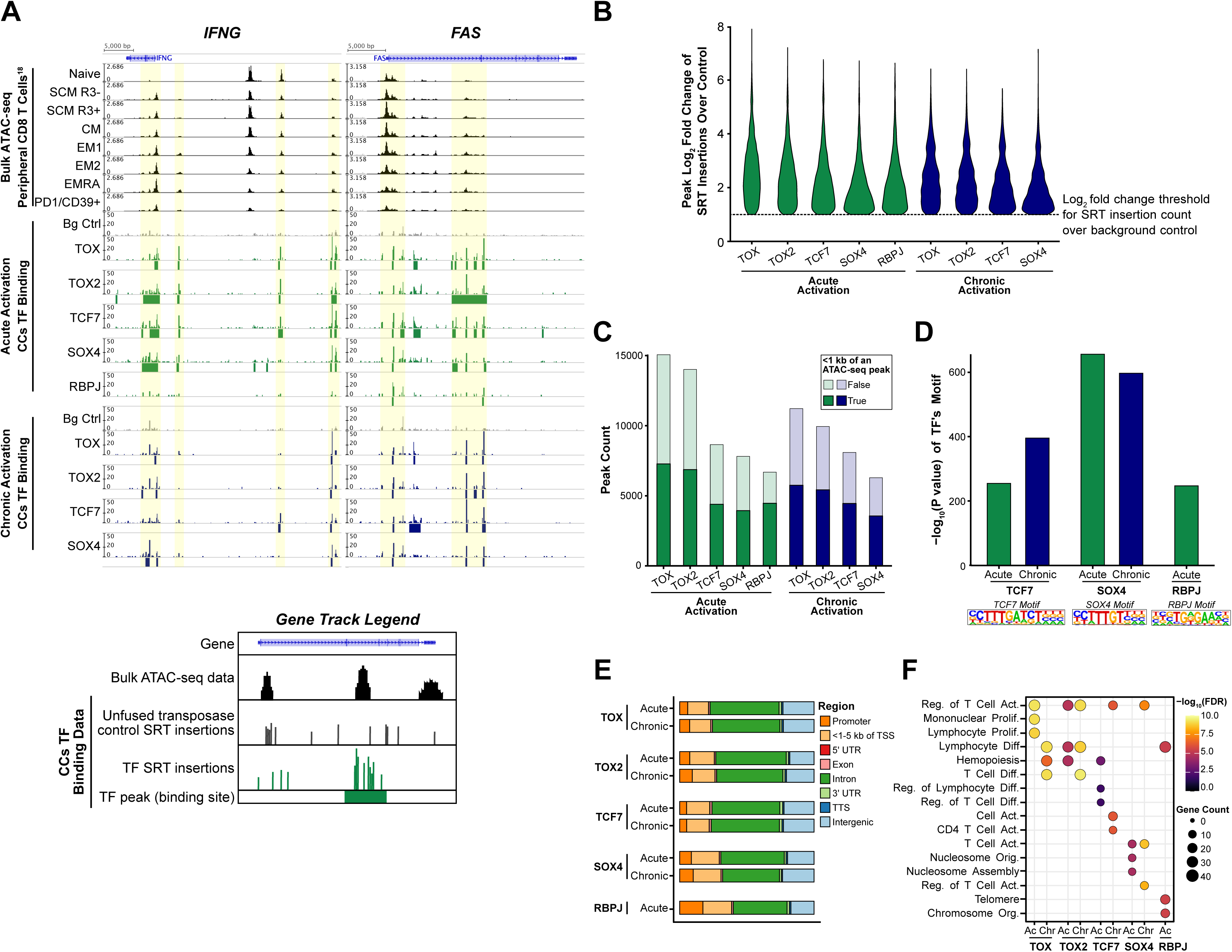
The Calling Cards assay robustly maps transcription factor binding sites in primary human CD8 T cells. **(A)** Normalized SRT insertion counts at the *IFNG* (*left*) and *FAS* (*right*) loci. TF peaks are indicated by the bars under the SRT insertion count tracks. The background control (Bg Ctrl) is the unfused HyPBase transposase. Example regions are highlighted, and bulk ATAC-seq profiles of peripheral human CD8 T cell subsets are included as a reference. The subsets include naïve, CXCR3-stem cell memory (SCM R3-), CXCR3+ SCM (SCM R3+), central memory (CM), CD27+ effector memory (EM1), CD27-effector memory (EM2), effector memory re-expressing CD45RA (EMRA), and exhausted (PD1/CD39+). **(B)** Violin plot of the log_2_ fold change of normalized SRT insertions over the unfused HyPBase background control in the defined peaks. A dashed line at 1 indicates the log_2_ fold change threshold for defining a peak. A minimum of ten normalized SRT insertions for the TF was also used as a threshold. **(C)** Number of defined peaks for each TF in the acute and chronic activation conditions colored by whether the peak is within 1 kb of a bulk ATAC-seq peak in the peripheral human CD8 T cell dataset^18^. **(D)** -log_10_(P value) of HOMER motif enrichment analysis of the TCF7, SOX4, and RBPJ motifs within the defined peaks of the corresponding TF. **(E)** Proportion of peaks within genomic regions. **(F)** Top three enriched GO processes for the genes bound within 1 kb of the transcription start site by each TF.

We next conducted several analyses to validate the accuracy of the binding sites. First, although these TFs were mapped *in vitro*, we found considerable overlap in TF binding sites with *in vivo* accessible chromatin regions. Approximately 50-60% of the TF peaks were within one kilobase of a highly statistically significant ATAC-seq peak (ATAC-seq peaks filtered by q-value<0.001 in >1 sample) from a published dataset of *in vivo* peripheral human CD8 T cell subsets (**Fig. 2C**)^18^. TF peaks that were not near an ATAC-seq peak may represent sites where TFs open the chromatin in a pioneer-like fashion, or they may be chromatin regions that are accessible only upon acute or chronic activation. We next confirmed that the TCF7, SOX4, and RBPJ canonical motif sequences were significantly enriched in the respective TF’s binding sites (**Fig. 2D**; HOMER motif enrichment analysis, -log_10_(P value)>200 for all TFs). TOX and TOX2 were excluded from this analysis as neither binds DNA in a sequence-specific manner, and both lack known motifs^19^. Finally, we validated that the binding sites for each condition were significantly enriched within five kilobases of transcription start sites (**Fig. 2E**; ∼25% of binding sites; one-sided permutation test, n=1,000, p<1×10^−3^ for all conditions). The genes bound within one kilobase of the transcription start site also had the expected enrichment of T cell-related gene ontology (GO) processes (**Fig. 2F**). Collectively, these data demonstrate that CCs accurately maps TF binding sites in primary human CD8 T cells, providing a robust list of putative direct targets of TOX, TOX2, TCF7, SOX4, and RBPJ.

### Calling Cards transcription factor binding data identifies the convergence of TCF7 and TOX binding at enhancers of peripheral human memory CD8 T cells

We next sought to utilize the TF binding data to gain insights into the biological roles of these TFs, specifically the complex interplay of the TOX and TCF7 regulatory networks. Both TOX and TCF7 influence gene expression through transcriptional control and modification of chromatin histone acetylation, which is important in the regulation of enhancers^20–22^. In mice, the interaction of the TOX and TCF7 networks is important specifically in exhaustion, where TOX is hypothesized to prevent activation-induced cell death, thereby allowing the TCF7-regulated stem-like exhausted population to form^23^. However, in human CD8 T cells, TOX and TCF7 are co-expressed in both memory and exhaustion, suggesting a possible functional role of these TFs in human memory phenotypes^24^. These CD8 T cell memory phenotypes differentiate from naïve CD8 T cells and have been proposed to follow a trajectory from naïve to stem cell memory (SCM), central memory (CM), effector memory (EM), and lastly, the terminal EM re-expressing CD45RA (EMRA) state^18,25^. As in the differentiation from stem-like exhaustion to terminal exhaustion, the trajectory to the EMRA phenotype corresponds with decreasing expression of TCF7 and stemness genes alongside increasing expression of TOX and effector genes^25^. Thus, because both TCF7 and TOX influence histone acetylation, which is central to enhancer regulation, we hypothesized that these two TFs might bind enhancer regions of stemness and effector genes that are differentially accessible in memory CD8 T cell phenotypes.

To investigate this possibility, we integrated the acute activation TCF7 and TOX CCs binding data with a dataset of 106 paired bulk RNA- and ATAC-seq samples of peripheral naïve and memory subsets of human CD8 T cells^18^. First, we confirmed that TOX mRNA was expressed in human memory CD8 T cells with relatively increased expression in the EM and EMRA populations, in agreement with prior studies (**Fig. 3A**)^24^. In support of a potential role of TOX in memory, the EM and EMRA subsets expressed TOX mRNA to a similar extent as the peripheral exhausted subset in this dataset (**Fig. 3A**). We next used the ENCODE-rE2G database of human CD8 T cell enhancers, which were defined by chromatin marks and 3D chromatin contacts, to annotate enhancer-associated ATAC-seq peaks and link them to genes (**Supplementary Table 2**)^26^. We refer to ATAC-seq peaks and their corresponding genes as enhancer-gene pairs. Of the 92,767 total ATAC-seq peaks, 62,088 (67.9%) overlapped with an annotated enhancer, and we linked an additional 2,061 (2.2%) ATAC-seq peaks within five kilobases of a transcription start site to their proximal genes. This amounted to 151,141 enhancer-gene pairs, which we then further filtered to the TOX- or TCF7-bound enhancer-gene pairs that were dynamically co-regulated across the memory subsets. First, we identified 14,872 bound enhancer-gene pairs (5,734 ATAC-seq peaks), of which TOX bound 12,069 pairs (4,505 ATAC-seq peaks) and TCF7 bound 7,146 pairs (2,856 ATAC-seq peaks; **Supplementary Table 3**). We next subset these to a final set of 1,481 bound and dynamically co-regulated enhancer-gene pairs (1,258 ATAC-seq peaks) based on a significant positive association between the enhancer accessibility and gene expression (linear regression FDR<0.05, β>0.15, R^2^>0.15 and Spearman correlation FDR<0.05). Suggesting a specific regulatory role at enhancers of memory subsets, TOX and TCF7 binding sites were enriched at this final set of enhancer regions relative to all ATAC-seq peaks (one-sided permutation test, n=1,000, p<1×10^−3^ for both TFs). Additionally, TOX and TCF7 co-bound 34% of these enhancer regions, representing a significant degree of overlap (**Fig. 3B**; Fisher’s exact test, p<2.2×10^−16^). Together, these data indicate a surprisingly high degree of enrichment and overlap of TCF7 and TOX binding at differentially accessible enhancer regions of memory CD8 T cells.

**Figure 3.**
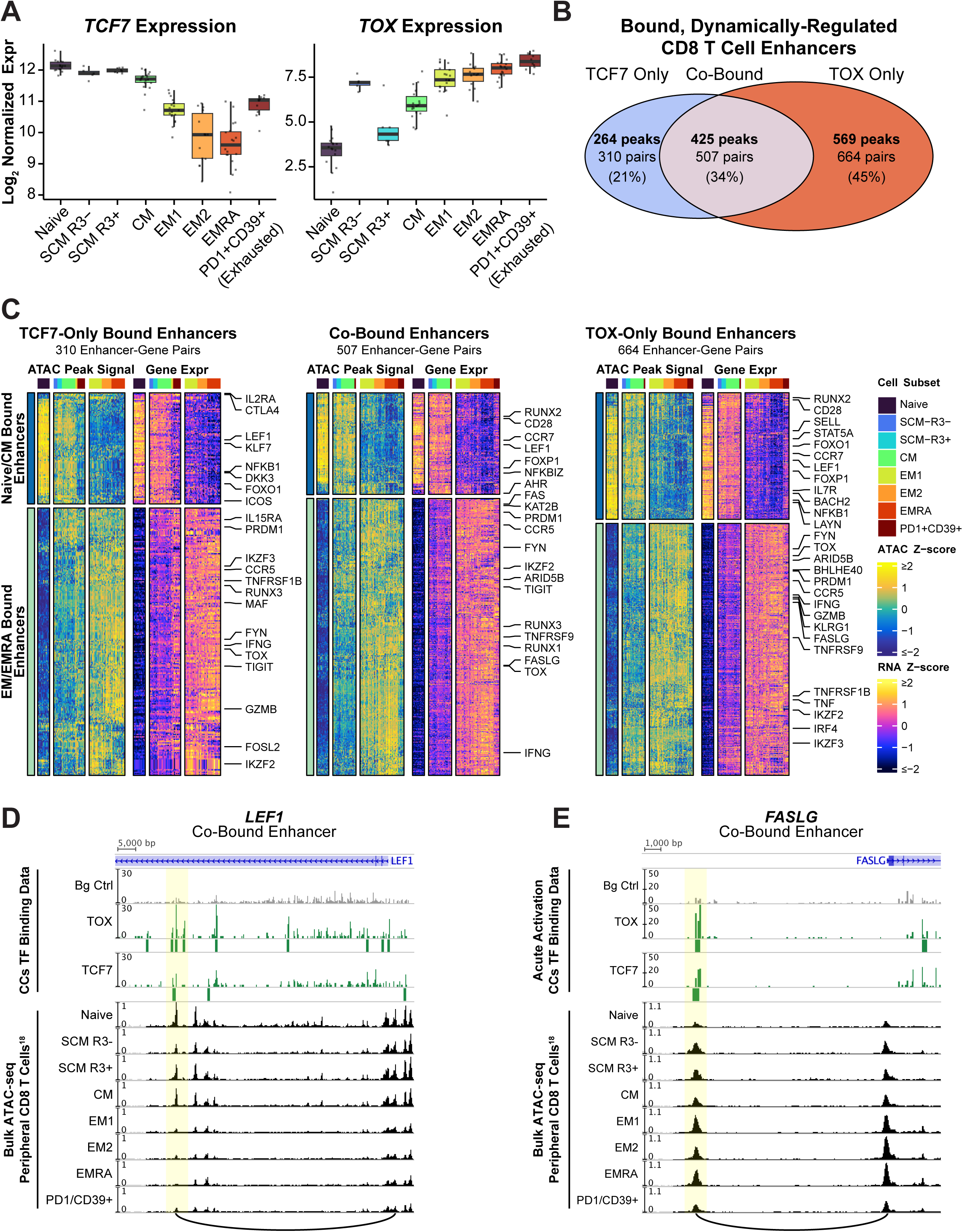
TCF7 and TOX converge on dynamically regulated enhancer regions of peripheral human CD8 T cells. **(A)** Log_2_ normalized expression (Expr) of TCF7 and TOX in bulk RNA-seq data of human peripheral CD8 T cell subsets, including naïve, CXCR3-stem cell memory (SCM R3-), CXCR3+ SCM (SCM R3+), central memory (CM), CD27+ effector memory (EM1), CD27-effector memory (EM2), effector memory re-expressing CD45RA (EMRA), and exhausted (PD1+CD39+). **(B)** Venn diagram of dynamically regulated enhancer-associated ATAC-seq peaks and the corresponding enhancer-gene pairs bound by TCF7 only, TOX only, or co-bound. **(C)** Heatmaps of enhancer-gene pairs showing the enhancer region ATAC-seq peak signal (labeled as ATAC) and the expression of its corresponding gene in the RNA-seq data (labeled as RNA) paired by row. These data are plotted for the enhancers bound only by TCF7 (*left*), co-bound (*middle*), or bound only by TOX (*right*). The cell subset is annotated at the top of the heatmap, and the broad classification of the enhancers (Naïve/CM or EM/EMRA) is annotated on the left. **(D-E)** Example gene tracks of the co-bound enhancer regions (*highlighted*) for *LEF1* (**D**) and *FASLG* (**E**) including the normalized SRT insertion counts for TOX, TCF7, and the unfused HyPBase background control (Bg Ctrl). The TF peaks, or binding sites, are indicated by the bars under the SRT insertion count tracks.

We next analyzed the genes of the bound enhancer-gene pairs. We plotted the ATAC-seq peak signal and gene expression for each enhancer-gene combination paired by row in adjacent heatmaps for the TCF7-only, TOX-only, and co-bound enhancers (**Fig. 3C**). Through k-means clustering, we identified two categories of enhancer-gene pairs, one associated with naïve and CM subsets and the other related to EM and EMRA subsets (**Fig. 3C**). Among the naïve/CM-associated enhancers, TCF7 and TOX co-bound to the enhancer regions of genes with critical roles in memory or stemness (**Fig. 3C**, *top half of the co-bound enhancers heatmap*), including *LEF1*^27^ (plotted in **Fig. 3D**), *FOXP1*^28^, and *RUNX2*^29,30^. However, most of the TCF7-only, TOX-only, and co-bound enhancers were associated with genes expressed in the EM/EMRA subsets, which were ∼65-75% of each set (**Fig. 3C**, *bottom half of each heatmap*). The co-bound EM/EMRA-associated enhancer-gene pairs (**Fig. 3C**, *bottom half of the co-bound enhancers heatmap)* featured key effector-related genes, including *FASLG*^31^ (plotted in **Fig. 3E**), *RUNX3*^32^, and *PRDM1*^33^. This convergence of TCF7 and TOX binding at enhancers of stemness and effector genes in human CD8 T cells is similar to their role in mouse CD8 T cell exhaustion, where they have been found to regulate an axis of memory- and effector-related genes^23,34,35^. Due to their involvement in histone acetylation, TCF7 and TOX may have unique synergistic or competing effects on chromatin accessibility at the co-bound enhancer regions that may not be involved at the enhancers bound by only TCF7 or TOX^20–22^. However, the precise combinatorial effects of TCF7 and TOX at these co-bound enhancers remain unclear. While TCF7 is known to possess intrinsic histone deacetylase activity, TOX has only been shown to *interact* with histone acetyltransferases, and the effects of this interaction are not well studied^20^. Collectively, we identified convergence of TCF7 and TOX binding at dynamically co-regulated enhancer-gene pairs of stemness and effector functions, suggesting an unappreciated functional role for these two TFs in memory subsets.

### Integration of transcription factor binding data with single cell RNA sequencing data identifies co-bound gene expression programs corresponding to fundamental CD8 T cell states

In addition to our data supporting TOX and TCF7 involvement in memory states, these TFs have a canonical role in exhaustion^36^. However, it has yet to be determined whether these TFs act upon a common gene program that is expressed in both memory and exhaustion states or regulate subset-specific programs. We therefore next aimed to define the gene programs that may be targeted by TOX and TCF7 in both memory and exhaustion, hypothesizing they would bind genes of subset-specific programs due to the functional differences between these states^36^. Additionally, to expand our TF analysis beyond TOX and TCF7, we also mapped three other major TFs with important roles that lack defined binding sites—TOX2, SOX4, and RBPJ^37–41^. For this analysis, we developed an analytical framework suited for scRNA-seq data to identify gene programs rooted in empiric evidence of TF binding (**Supplementary Fig. 5**). In this approach, we first identified all genes bound at an ENCODE-rE2G enhancer^26^ among the co-regulated enhancer-gene pairs or a promoter using the acute and chronic activation CCs TF binding data to capture the full breadth of potential targets (**Supplementary Table 3**). Next, using a published scRNA-seq atlas of human CD8 T cells that includes both memory and exhaustion states, we determined the association between TF expression and each of its bound genes (**Fig. 4A; Supplementary Table 4**)^42^. We then subset these genes to only those that had a positive association with TF expression to obtain the final TF-bound and associated gene sets (**Supplementary Table 5**). This gene set represents putative positively regulated targets or targets involved in negative feedback where TF expression increases with increasing gene expression. Lastly, we quantified the expression of each TF and calculated the module score of its bound and associated gene set in the scRNA-seq atlas (**Fig. 4B**). This illustrated that the expression of the TF-bound genes spanned subsets, such as TOX binding to genes expressed in both memory and exhaustion subsets (**Fig. 4B**). Additionally, the expression of each TF (**Fig. 4B**, *top*) was not tightly correlated with the module score of its bound and associated gene set (**Fig. 4B**, *bottom*), indicating that subset-specific co-factors or epigenetic profiles likely determine the ultimate effect of the TF on its bound genes. Thus, the variation in the module scores of the TF-bound and associated gene sets likely represents expression of subset-specific, distinct gene programs rather than differential expression of the full gene set. In other words, TOX may bind and regulate distinct gene programs in memory and exhaustion subsets rather than the same program in both subsets. Correspondingly, different TFs may converge on subset-specific programs.

**Figure 4.**
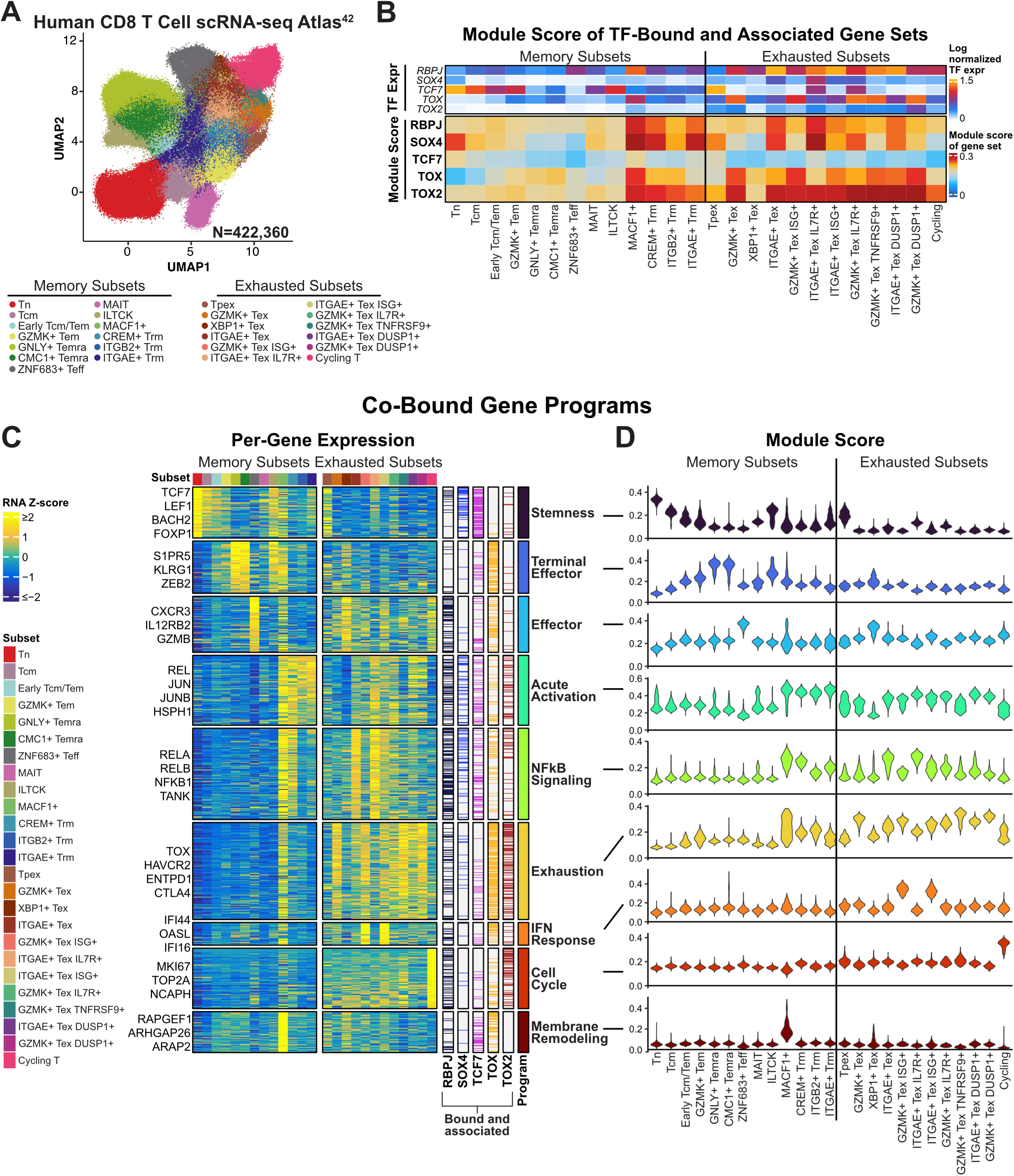
Calling Cards transcription factor binding data reveals co-bound gene expression programs in single cell RNA-sequencing data. **(A)** UMAP plot of the scRNA-seq atlas of human CD8 T cells (N = 422,360 cells)^42^. **(B)** Average log-normalized TF expression (*top*) and module score of the TF-bound and positively associated gene sets (*bottom*) per scRNA-seq subset among metacells (*k*=15). **(C)** Heatmap of the Z-scored average RNA expression of each gene within the co-bound gene programs defined by NMF among the metacells. The marker genes of each program are annotated on the left, and on the right, each TF’s bound and positively associated genes are indicated with a horizontal line at the position of each gene. The name of the gene program is annotated on the far right. **(D)** Violin plots of the module score for each co-bound gene program listed in the same order as (**C**) among the cell subsets.

To investigate co-bound gene programs, we used non-negative matrix factorization (NMF) with all unique genes of the TF-bound and positively associated gene sets as the input (**Supplementary Fig. 5**). We defined nine co-bound gene programs corresponding to fundamental CD8 T cell states or functions and assigned names based on marker genes (**Supplementary Table 6**). Specifically, we identified programs related to stemness (*LEF1*, *BACH2*), terminal effector differentiation (*KLRG1*, *ZEB2*), the effector state (*IL12RB2*, *GZMB*), acute activation (*REL*, *JUN*), NF-κB signaling (*RELA, RELB*), exhaustion (*TOX*, *ENTPD1*), interferon response (*IFI44*, *OASL*), cell cycle (*MKI67*, *TOP2A*), and membrane remodeling (*RAPGEF1*, *ARAP2*) (**Fig. 4C**). We quantified the expression of the co-bound programs on a per-gene level and marked the bound and positively associated genes of each TF to visualize the proportion of TF binding in the programs (**Fig. 4C**; **Supplementary Table 7**). As a metric of overall program expression, we also plotted the module score of the program genes in each scRNA-seq subset (**Fig. 4D**). This comprehensive analysis revealed TOX binding to 82% of the exhaustion gene program, representing genes that TOX directly regulates to mediate its hypothesized role as a master regulator of exhaustion^20,23,34,35,43,44^ (**Fig. 4C, D**). Additionally, consistent with its function in promoting self-renewal, TCF7 bound 80% of the stemness program genes, which were highly expressed in the memory and stem-like progenitor exhausted (Tpex) subsets, as expected^45–47^ (**Fig. 4C, D**). We also observed an interaction between the TOX and TCF7 regulatory networks in exhaustion: relative to the Tpex subset, the TCF7-bound stemness program was *decreased*, while the TOX-bound exhaustion program was *increased* in the differentiated exhaustion subsets (**Fig. 4C, D**). Mirroring our prior analysis (**Fig. 3**), TOX also bound the terminal effector program (76% of genes bound), which was largely exclusive to the memory subsets and enriched in the EMRA subset (**Fig. 4C, D**). Interestingly, like its relation to the exhaustion program, the TCF7-bound stemness program was anti-correlated with the TOX-bound terminal effector program, substantiating a potential interplay between the functions of TCF7 and TOX in memory subsets (**Fig. 4C, D**). Overall, these data indicate that TOX and TCF7 coordinate related gene programs in *both* memory and exhaustion, where TCF7 regulates stemness, and depending on the cell state, TOX acts upon an opposing exhaustion or terminal effector program.

In addition to TOX and TCF7, we identified TOX2 binding to genes consistent with its proposed functions, including occupancy at 55% of the exhaustion program genes^41^ and 50% of cell cycle genes^40^ (**Fig. 4C; Supplementary Table 7**). While SOX4 and RBPJ are not as well studied, we gained insights into their roles with our analysis, where we found SOX4 bound to stemness-related genes (47% bound). This program may underlie its unknown function in the SOX4+ recent thymic emigrant naïve CD8 T cell subset^38^. By contrast, RBPJ bound to a high proportion of genes in programs broadly related to T cell activation, including cell cycle (45%), NF-κB signaling (59%), acute activation (48%), and effector (49%) programs (**Fig. 4C**). Interestingly, the acute activation and NF-κB signaling gene programs were generally co-expressed with the exhaustion program, which arises in response to persistent activation (**Fig. 4C, D**)^20^. However, these programs were predominately bound by distinct TFs, with TOX binding most exhaustion genes and RBPJ binding the acute activation and NF-κB signaling genes (**Fig. 4C; Supplementary Table 7**). Thus, these programs are likely differentially regulated through the actions of unique TFs, which may underlie the distinct gene expression changes that occur in response to the chronicity of T cell activation. Within this framework, RBPJ regulates genes fundamentally important in activation, such as *RELA* and *NFKB1.* Then, as TOX increases with chronic activation, a known phenomenon^20,34^, it may promote a coinciding exhaustion program of inhibitory genes (e.g., *CTLA4*, *ENTPD1*) that hinders activation-induced cell death. However, while most TFs tend to be ascribed a single function, such as TOX in exhaustion, our data cautions against this interpretation, instead illustrating that each TF binds genes of multiple programs (**Fig. 4C**). Together, our analytical framework to integrate CCs TF binding data with scRNA-seq data improves upon unsupervised program analyses by defining gene programs that are now rooted in empiric TF binding data. These programs therefore more closely resemble true coordinated transcriptional programs and highlight the value of TF binding data in disentangling cell states.

### Development of TFlex—a scalable, transposon-based method for multiplexed mapping of transcription factors

The integrative analysis of CCs TF binding data with the multi-omic bulk sequencing dataset and the scRNA-seq atlas led to key biological insights, some of which suggested alternatives to the leading hypotheses on TF function, as in the case of TOX. We therefore sought to further scale our TF analysis platform for primary human T cells by modifying the standard CCs method to create ***TFlex***: an antibody-independent, transposon-based method uniquely suited for multiplexed mapping of paralogous natural and synthetic TFs (**Fig. 5A**). In standard CCs, only 25 barcodes can be incorporated because these are within the terminal repeat sequence and affect HyPBase transposition efficiency but are crucial for detecting independent insertion events^48^. We therefore redesigned the SRT in TFlex to position the barcodes upstream of the terminal repeat sequence (**Fig. 5A**). This allowed for incorporation of both unlimited SRT barcodes, and importantly, an independent sample barcode. This sample barcode permits multiplexed TF mapping as the SRT containing the sample barcode is stably integrated into each cell’s genomic DNA by HyPBase in the TF-HyPBase fusion protein (**Fig. 5A**). However, repositioning the barcode region required that the Read 1 sequence started upstream of the terminal repeat sequence, which is 306 base pairs (**Fig. 5A**). Consequently, the Read 1 sequence could no longer be used to map the junction of the SRT and genomic DNA from standard short-read sequencing. Therefore, we devised a strategy to ascertain the sample and SRT barcodes from Read 1 and map the TF binding sites from the paired Read 2, which begins on genomic DNA and moves towards the terminal repeat sequence of the SRT (see **Methods**; **Fig. 5A**). In practice, T cells can be electroporated with separate pools of TFlex SRT vectors each containing a unique sample barcode and diverse SRT barcodes (**Fig. 5B**). Immediately after electroporation, T cells can be combined into a single culture and RNA-based library preparation followed by demultiplexing samples in the sequencing data on the basis of the sample barcode (**Fig. 5A, B**).

**Figure 5.**
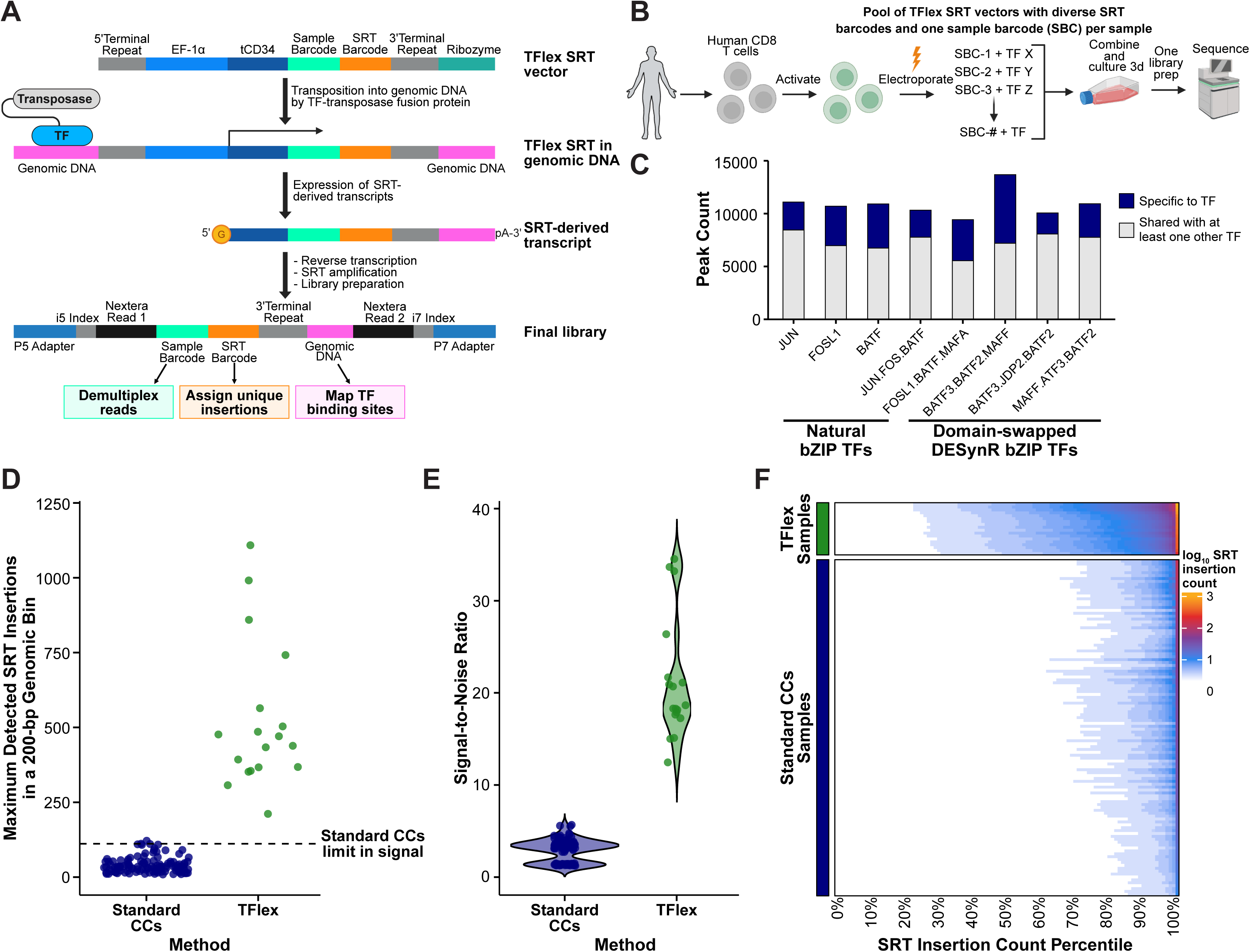
TFlex self-reporting transposons enable multiplexed TF mapping with improved dynamic range relative to standard Calling Cards. **(A)** Schematic of the TFlex SRT vector and library structure. The sample and SRT barcodes are positioned upstream of the 3’ terminal repeat sequence. Upon integration of the SRT into genomic DNA, transcripts are produced that contain the sample barcode, SRT barcode, 3’ terminal repeat, and downstream genomic DNA, which are detected in the library sequencing data. **(B)** Schematic of TFlex workflow. Each sample is electroporated with a pool of TFlex SRT vectors containing diverse SRT barcodes and a specific sample barcode (SBC). Because the SRT becomes integrated into the genomic DNA, cells can be combined immediately after electroporation. **(C)** Peak counts for the three natural bZIP TFs (JUN, FOSL1, and BATF) and five domain-swapped DESynR bZIP TFs mapped in a multiplexed manner with TFlex. The domain-swapped TF names refer to the N-terminal domain, DNA-binding domain, and C-terminal domain in that order and separated by a period. **(D)** Maximum detected SRT insertions in a 200-bp genomic bin for standard CCs and TFlex samples. **(E)** Signal-to-noise ratio of standard CCs and TFlex samples defined as the ratio of the mean SRT insertion counts in the top 20^th^ percentile of 200-bp genomic bins to that of the bottom 20^th^ percentile. **(F)** Heatmap of the log_10_ SRT insertion counts in 200-bp genomic bins ordered by percentile rank for standard CCs and TFlex samples.

To validate the multiplexed capacity of TFlex, we simultaneously mapped eight paralogous bZIP family TFs. These included JUN, FOSL1, BATF, and five domain-swapped DESynR TFs previously found to enhance the anti-tumor functions of human T cells^13^. We identified approximately 10,000 peaks for each TF, defined as a log_2_ fold change of normalized SRT insertions greater than or equal to 1 relative to the unfused HyPBase background control and at least eight normalized SRT insertions (see **Methods**; **Fig. 5C; Supplementary Table 1**). Demonstrating the fidelity of TFlex to define unique binding sites in multiplexed samples, we identified peaks specific to each TF relative to all of the other seven TFs (**Fig. 5C**). However, as expected because all of these TFs possess a bZIP DNA-binding domain, most binding sites were shared with at least one of the other seven TFs (**Fig. 5C**). Indeed, the bZIP motif sequence was significantly enriched among the peaks of all natural and domain-swapped DESynR bZIP TFs (**Supplementary Fig. 6**; HOMER motif enrichment analysis, -log_10_(P value)>250 for all). We next compared TFlex to standard CCs using our data from primary human CD8 T cells. By removing the limit in SRT barcodes in the TFlex method, the maximum detected SRT insertions in a 200-base pair genomic bin increased approximately 10-fold, from 122 in standard CCs to 1,112 in TFlex (**Fig. 5D**). The mean signal-to-noise ratio was also increased approximately 10-fold, which was 2.5 in standard CCs relative to 21.2 in TFlex (**Fig. 5E**). Lastly, to visualize the dynamic range of each method, we plotted the SRT insertion count per 200-base pair genomic bin ordered by percentile rank (**Fig. 5F**). While the majority of data in standard CCs was within the top 80^th^ to 100^th^ percentiles, the dynamic range of TFlex was substantially greater, with considerable SRT insertion counts spanning the 20^th^ to 100^th^ percentiles (**Fig. 5F**). Ultimately, these data highlight TFlex as a scalable TF mapping method with substantial dynamic range in TF binding signal that is capable of multiplexed TF mapping in primary human T cells.

### Synthetic domain-swapped bZIP transcription factors possess unique emergent properties relative to the parent TFs of their constituent domains

We next sought to use our TFlex data to investigate the properties of the JUN.FOS.BATF DESynR TF, which was engineered from the N-terminal domain of JUN, DNA-binding domain of FOS, and C-terminal domain of BATF (**Fig. 6A**)^13^. Although created from natural TF domains, JUN.FOS.BATF was found to imbue primary human T cells with enhanced anti-tumor features relative to its natural TF counterparts^13^. This may be due to either JUN.FOS.BATF acquiring distinct binding sites at genes that regulate T cell function or recruitment of unique co-factors at shared binding sites that differentially regulate target gene expression.

**Figure 6.**
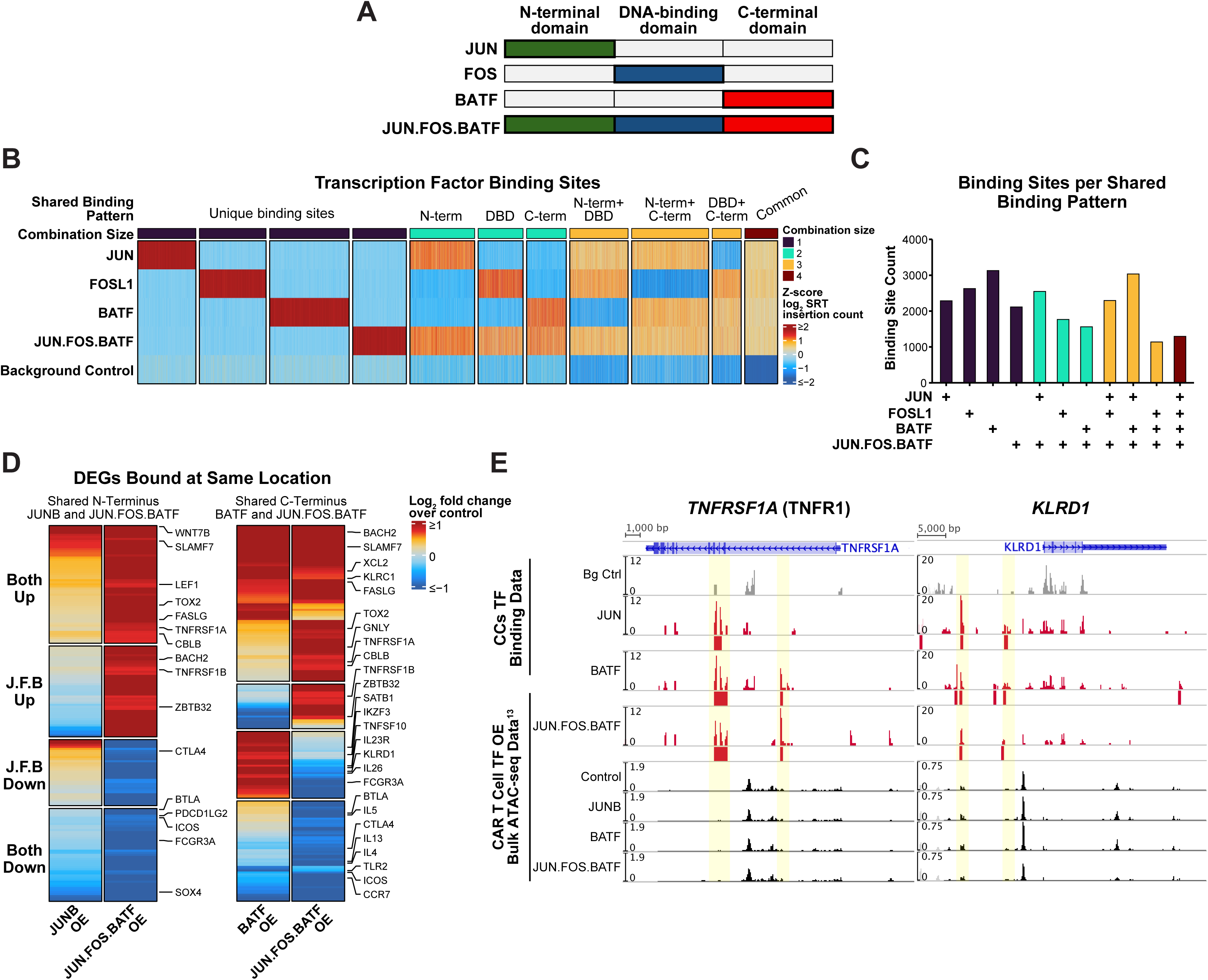
TFlex reveals distinct DNA binding and target gene effects of natural and domain-swapped bZIP family transcription factors. **(A)** Schematic of the JUN.FOS.BATF domain-swapped DESynR TF illustrating that it contains the N-terminal domain of JUN, DNA-binding domain of FOS, and C-terminal domain of BATF. **(B)** Heatmap of Z-scored log_2_ SRT insertion counts at TF binding sites that were manually split based on the shared binding pattern. Each column is a peak, and each row is the Z-scored log_2_ SRT insertion count for the TF in each peak. For visualization, overlapping peaks were merged into a single peak such that each column is a unique genomic region. The background control is the unfused HyPBase. **(C)** Peak counts for each shared binding pattern. **(D)** Log_2_ fold change of gene expression for JUNB, BATF, and JUN.FOS.BATF overexpression relative to control from pseudobulked scRNA-seq data^13^ of genes bound at the same location by JUN and JUN.FOS.BATF (*left*) or BATF and JUN.FOS.BATF (*right*). The general pattern of expression, such as JUN.FOS.BATF uniquely up-regulated (J.F.B. up), is annotated on the left. **(E)** Gene tracks illustrating shared binding sites (*highlighted*) at genes with differential expression relative to control between JUN.FOS.BATF and a natural TF. Bulk ATAC-seq data from CAR T cells for each TF overexpression (OE) condition are shown as a reference^13^.

To directly test these possibilities, we first analyzed the overlap of JUN.FOS.BATF binding sites with each of its parent TFs. We expected that all JUN.FOS.BATF binding sites would be shared with at least one of its parent TFs based on the shared domains. Instead, we found that approximately 15% of JUN.FOS.BATF binding sites were unique despite JUN.FOS.BATF having binding sites similarly enriched for the bZIP motif as its parent TFs (**Fig. 6B; Supplementary Fig. 6**). Based on the central hypothesis that the DNA-binding domain determines binding site selection, we expected that the shared binding sites of JUN.FOS.BATF would be biased towards FOSL1 or JUN, which have similar DNA-binding domains to FOS^49^. However, we did not observe any bias of JUN.FOS.BATF binding site overlap towards a specific parent TF (**Fig. 6C**). These data illustrate that domain-swapped TFs can acquire unique DNA binding site selection properties that are not explained by the DNA-binding domain, which may contribute to their distinct functional effects in T cells.

We next investigated whether JUN.FOS.BATF differentially regulated target gene expression at binding sites that were shared with a parent TF. Using RNA-seq data from JUNB-, BATF-, and JUN.FOS.BATF-overexpressing chimeric antigen receptor (CAR) T cells^13^, we first defined the differentially expressed genes relative to control for each condition. We then subset the genes to those that were bound by both JUN.FOS.BATF and JUN or BATF and differentially expressed in either condition. When comparing the co-bound genes of JUN.FOS.BATF and JUN, the parent TF of the N-terminal domain, approximately 40% of these genes had discordant expression, where the gene was decreased by JUNB yet increased by JUN.FOS.BATF and vice versa (**Fig. 6D**, *left*). The remaining genes were relatively concordant, suggesting a shared effect on gene expression that may be mediated through the N-terminal domain although the extent of differential expression varied. We observed a similar phenomenon for genes co-bound by BATF and JUN.FOS.BATF, which share the C-terminal domain (**Fig. 6D**, *right*). Approximately 30% of these genes were discordant with the remainder being either increased or decreased by BATF and JUN.FOS.BATF overexpression relative to control. While we did not identify a broader program of the genes differentially regulated by JUN.FOS.BATF, there were several genes with established, important roles in human T cells: *TNFRSF1A* (plotted in **Fig. 6E**, *left*), *KLRD1* (plotted in **Fig. 6E**, *right), TNFRSF1B, LEF1*, *BACH2*, *PDCD1LG2*, *CTLA4*, and *ICOS*. Thus, neither the target gene effects nor DNA binding sites of domain-swapped TFs can be estimated as the sum of their domains. Instead, empiric, experimentally-driven DNA binding data is required to determine the unique emergent properties of these synthetic TFs that drive their DNA binding and functional effects on target genes. This has important implications in refining the structure of synthetic TFs to program T cells into favorable cell states for tumor elimination or other therapeutic uses. Collectively, our data highlight the importance of scalable TF mapping in primary human T cells to elucidate how TFs govern cell states and to prioritize therapeutic strategies, particularly within synthetic biology.

## DISCUSSION

TFlex is a major technological advancement that unlocks scalable, antibody-independent investigations of both natural and synthetic TFs in primary human T cells and other cell types. While chromatin immunoprecipitation-seq (ChIP-seq) has been adapted for mapping over 100 histone markers or TFs in parallel, it relies on specific anti-TF antibodies. These antibodies are typically unable to discriminate between paralogous TFs, such as TOX and TOX2, or between domain-swapped synthetic TFs, which often contain large regions from endogenous TFs. Furthermore, ChIP-seq requires chemical fixation and DNA fragmentation, which can introduce artifacts^10,50–52^. Other methods to map TFs, such as CUT&RUN, have had limited success for multiplexed TF mapping and would suffer from the same difficulties as ChIP-seq at discriminating between TFs with homologous regions^53–60^. While there are potential technical concerns with standard CCs or TFlex, including the fusion of the TF to HyPBase and exogenous expression, we have not found these to be major limitations. To address these concerns, we used an unfused HyPBase background control to remove binding sites driven by the transposase rather than the TF. We also validated that the TF binding sites possessed the expected TF motif and were localized to both the ATAC-seq peaks of *in vivo* CD8 T cells and the transcription start sites of T cell-related genes. Additionally, in prior studies, we have found that 90% of CCs peaks were within one kilobase of the peaks identified by ChIP-seq^14^, demonstrating the methods are generally highly concordant. Lastly, exogenous TF expression is not a limitation unique to standard CCs or TFlex as mapping TFs without a specific antibody in other methods would require expression of an epitope-tagged TF. Additionally, with CRISPR editing, it may be feasible to fuse the HyPBase to the TF in the genomic DNA such that the TF-HyPBase fusion protein is expressed at physiological levels. In summary, the potential limitations of TFlex are not likely to hinder key biological insights gained from multiplexed mapping of TFs.

Our analytical platform to integrate TF binding data with multi-omic sequencing to define TF-targeted gene programs is an improvement on existing computational algorithms that predict TF regulatory networks. Existing algorithms define TF binding *in silico* using canonical motif sequences^61^. However, motif presence is insufficient evidence for binding^62^, and a quarter of human TFs (∼400 total) do not have a known motif, including TOX and TOX2 and potentially many synthetic TFs^19,63–66^. TFlex now provides a way to feasibly obtain TF binding data at scale in primary human T cells, enabling an analysis of cell states and TF functions rooted in experimentally-determined DNA binding sites. With this approach, we identified gene programs related to activation that were co-expressed yet differentially bound by RBPJ and TOX, revealing distinct TF-mediated regulation of coinciding gene programs. Although the expression of these programs was similar in the scRNA-seq dataset, there are likely dynamic states of activation where these programs diverge. Beyond identifying transcriptional gene programs, TF binding data offers key insights into possible transcriptional and functional consequences of impaired TF function due to pharmacologic inhibition or mutations, for example. Mapping TFs may also guide hypotheses on the functional impacts of single-nucleotide polymorphisms (SNPs) at TF binding sites. Lastly, since gene expression is influenced by combinatorial roles of TFs, understanding which TFs bind and potentially regulate genes may be a prerequisite for thoroughly understanding transcriptional gene regulation. Collectively, our scalable TF analysis platform and its results add to the growing body of literature on capturing dynamic, combinatorial cell states and TF functions that more accurately reflect true biology compared to rigid definitions^42,67,68^.

By comprehensively profiling our TOX, TCF7, TOX2, SOX4, and RBPJ binding data, we gained key insights into putative TF-targeted gene programs that underlie known and potentially unrecognized functions, particularly for TOX and TCF7. Complementing the mouse studies indicating a role of TOX in exhaustion, our human CD8 T cell TOX binding data supports this TF as a major regulator of exhaustion and adds the critical piece of information of putative direct TOX targets^20,23,34,35^. However, in opposition to mouse models suggesting TOX is important only in exhaustion^20,23,34,35^, we instead found TOX binds a terminal effector program expressed in memory subsets in addition to binding an exhaustion program. Furthermore, our data supports that the TOX and TCF7 regulatory networks interact in *both* memory and exhaustion subsets, building upon the finding that TOX and TCF7 are co-expressed in both subsets of human CD8 T cells^24^. While the functions of TOX and TCF7 in exhausted mouse CD8 T cells are known to be cooperative^23^, our data provides new evidence that these TFs may interact through direct co-binding at enhancer regions. Our findings on TOX and TCF7 may be relevant to CAR T cell cancer therapies, which undergo a memory-to-exhaustion transition upon chronic stimulation that is poorly understood^37,69^. Given that these TFs bind to gene programs active in both memory and exhaustion states, they may be involved in this transition and be a potential node for therapeutic perturbation to improve T cell function.

While we have focused on TF-bound gene programs in primary human T cells, TFlex is broadly applicable to TF investigations due to its capacity for multiplexed, antibody-independent TF mapping. For example, because the SRT is stably integrated into the genome, TFlex can be used to longitudinally track TF binding during an experimental perturbation, such as activation of T cells or chemotherapy resistance of cancer cell lines. Additionally, as TFlex is RNA-seq based, it can be used with scRNA-seq for single-cell, multiplexed TF binding analyses. Lastly, we anticipate that TFlex will be central to studying domain-swapped or other synthetic TFs to deconstruct TF biology into the roles of specific domains, such as intrinsically disordered regions^70^. These insights on TF domains can support the rational development of synthetic TFs that imbue T cells with favorable therapeutic properties. In summary, we have developed a scalable method to map TFs, enabling discovery of gene networks underlying cell states and principles of TF function that could enable new avenues in therapeutic development.

## LIMITATIONS OF THE STUDY

Although we validated the accuracy of the TF binding sites, there are a few key limitations. First, in standard CCs or TFlex, the TF is fused to the HyPBase transposase, which may introduce false-positive or false-negative binding sites. Secondly, the TF-HyPBase fusion is overexpressed in the cell at levels that may be above the physiological expression. This may lead to an overrepresentation of binding sites that rarely occur at normal TF expression levels. Lastly, because the TF binding sites were determined *in vitro* in the context of acute or chronic activation, we cannot be certain that the binding and regulation of all putative gene targets occur *in vivo* in non-activation contexts. However, as addressed above, we have not found these limitations to hinder key biological insights gained from these methods.

## RESOURCE AVAILABILITY

### Lead Contact

Further information and requests for resources should be directed to the lead contact, Sidharth V. Puram.

### Materials Availability

The sequences of the SRT vectors and TF-HyPBase fusion proteins are supplied in **Supplementary Table 8**. Constructs generated for this manuscript are available by request to the lead contact.

### Data and Code Availability

The code used to process TFlex data, call TF peaks, analyze the data, and create supplementary tables is available in a GitHub repository (https://github.com/rileymullins/TFlex.git). The standard CCs and TFlex data has been deposited in a Synapse repository (syn69980172, https://doi.org/10.7303/SYN69980172). Raw FASTQ sequencing data will be made available on Gene Expression Omnibus. Publicly available ATAC-seq and RNA-seq data was obtained from the Gene Expression Omnibus: ATAC-seq and RNA-seq data of peripheral human CD8 T cells was obtained from GSE179613^18^, and ATAC-seq and RNA-seq data of CAR T cell natural and DESynR bZIP TF overexpression data was obtained from GSE280508 and GSE280509^13^.

## ACKNOWLEDGEMENTS

The authors thank Robert Schreiber, Takeshi Egawa, and Maxim Artyomov for insights guiding the analyses. We also thank the DNA Sequencing Innovation Lab of The Edison Family Center for Genome Sciences & Systems Biology at Washington University for assistance with the TFlex library design. We are grateful to GTAC at the McDonnell Genome Institute for sequencing services. This work was supported by K08CA237732 (NIH/NCI) (S.V.P.), R21DE031072 (NIH/NIDCR) (S.V.P.), R01GM123203 (NIH/NIGMS) (R.D.M.), R21DE031366 (NIH/NIDCR) (R.D.M., S.V.P.), R01DE032865 (NIH/NIDCR) (R.D.M., S.V.P.), P01CA245705 (NIH/NIGMS) (R.D.M.), and F31DE032562 (NIH/NIDCR) (R.D.Z.M.). Schematics were designed with Biorender.com.

## AUTHOR CONTRIBUTIONS

R.D.Z.M., R.D.M., and S.V.P. conceptualized the study. R.D.Z.M, R.D.M., and S.V.P. wrote and edited the manuscript with input from all authors. R.D.Z.M. and E.S. performed experiments. R.D.Z.M., J.Z., O.S., and R.S. created the computational pipeline to process the TFlex data. M.M. processed raw bulk ATAC-seq data for visualizations. R.D.Z.M. analyzed the data and prepared figures. R.D.Z.M., O.T., T.L.R., A.T.S., R.D.M. and S.V.P. guided experiments.

## SUPPLEMENTAL INFORMATION

**Supplementary Table 1.** Peaks called for each TF.

**Supplementary Table 2.** Processed CD8 T cell enhancer regions obtained from the ENCODE-rE2G database (Gschwind, R.A., et. al., *bioRxiv*, 2023).

**Supplementary Table 3.** Genes bound by the TFs at enhancer regions or transcription start site-proximal regions with simple linear regression and Spearman correlation statistics for the association between the enhancer region ATAC-seq peak and the corresponding gene expression. TF binding is indicated by a 1 for binding and a 0 for no binding.

**Supplementary Table 4.** Proportionality (*ρ_p_*) metric from the scRNA-seq atlas expression matrix of metacells (*k*=15) between each TF and the genes it binds at an enhancer region or ±1 kb of a transcription start site.

**Supplementary Table 5.** Final set of bound and negatively or positively associated gene targets of each TF that met the *ρ_p_* cutoff and were used in downstream analyses.

**Supplementary Table 6.** NMF-derived gene programs from all unique genes among the final bound and positively associated genes of TOX, TOX2, TCF7, SOX4, and RBPJ.

**Supplementary Table 7.** Proportion of each TF’s bound and positively associated genes within each NMF-derived gene program.

**Supplementary Table 8.** qPCR, standard CCs, and TFlex primer sequences; source and DNA sequences of TF-HyPBase fusion proteins; and DNA sequences of standard CCs and TFlex SRT-tCD34 minicircle vectors.

**Supplementary Figure 1.**
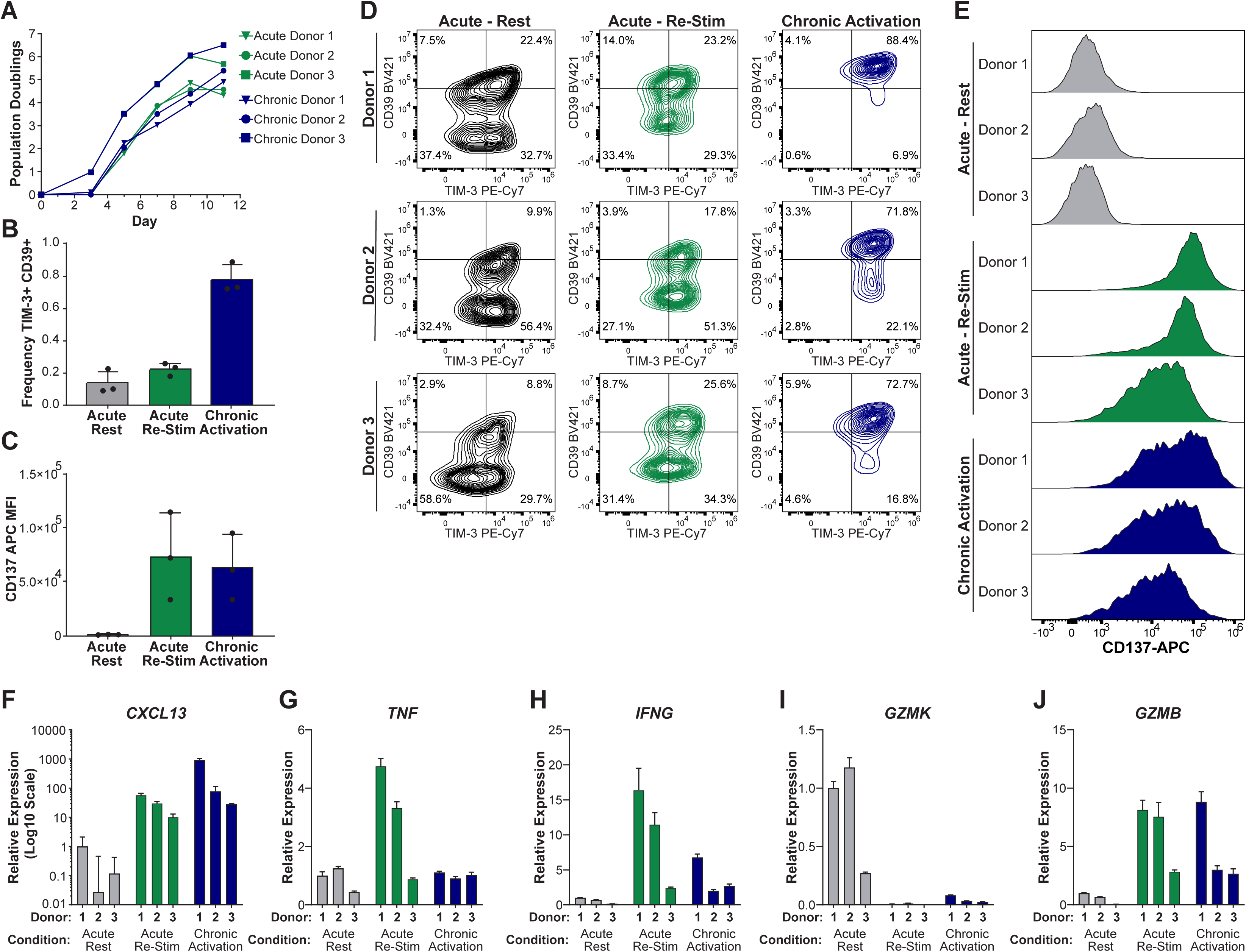
Characterization of primary human CD8 T cells subjected to *in vitro* acute or chronic activation. **(A)** Population doublings of CD8 T cells activated for 72 hours (acute) or 11 days (chronic) for each donor. **(B-J)** CD8 T cells were activated for 72 hours followed by 9 days of rest (Acute Rest), activated for 72 hours followed by 8 days of rest and then re-activated on day 11 for 24 hours (Acute Re-Stim), or activated for 12 days continuously (Chronic Activation). **(B)** Frequency of TIM-3+ CD39+ CD8 T cells. **(C)** Mean fluorescence intensity (MFI) of CD137. **(D)** Contour plots of TIM-3 and CD39 expression. **(E)** Histogram representations of CD137 expression normalized to the mode. **(F-J)** Relative gene expression plotted as 2^−ΔΔCt^ with the Donor 1 Acute Rest condition set as the reference. Error bars represent the range of expression values among technical triplicates for each gene.

**Supplementary Figure 2.**
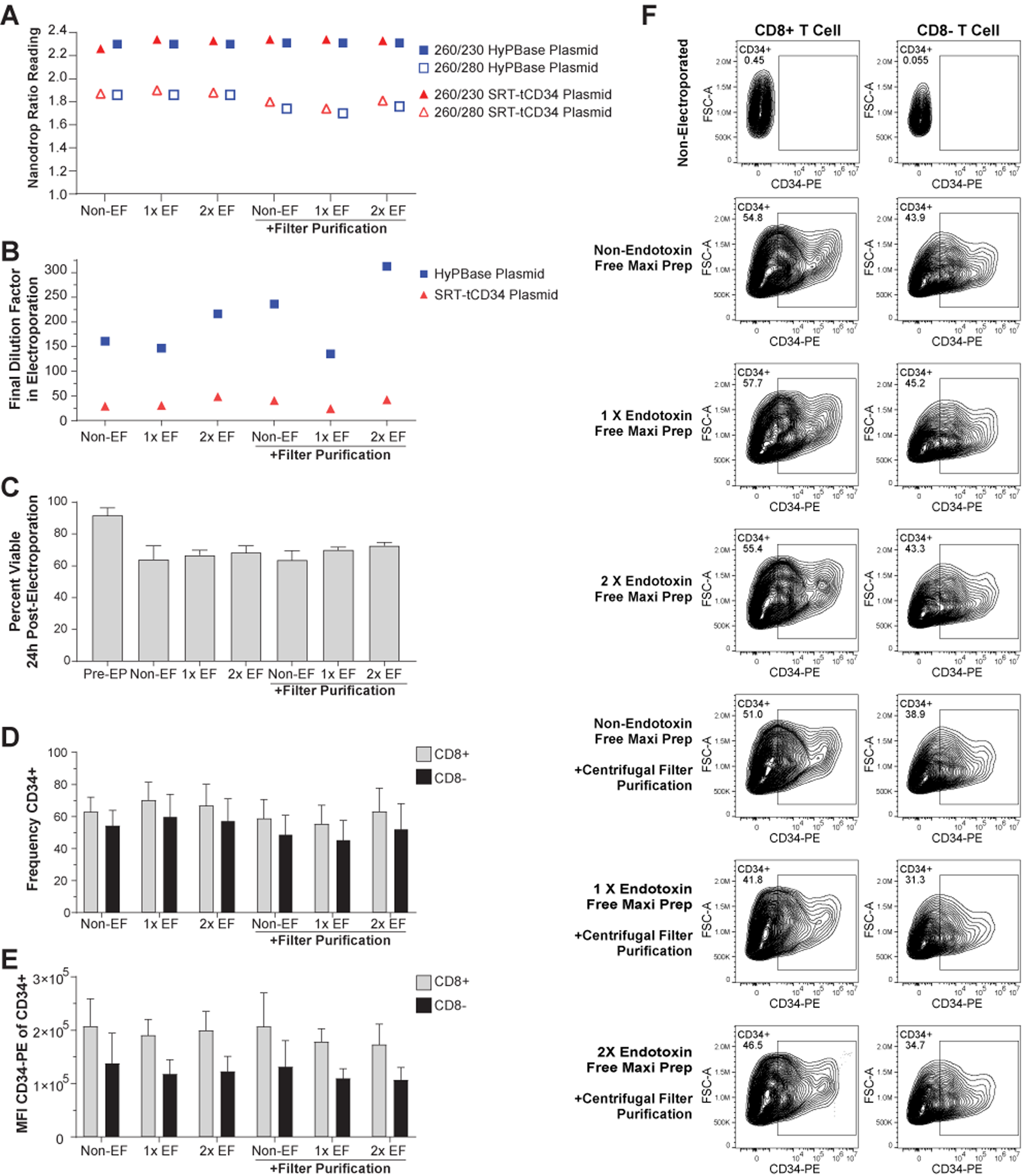
Comparison of HyPBase and SRT-tCD34 plasmid purification methods. **(A)** Ratios of 260/230 nm (filled blue square and filled red triangle) and 260/280 nm (open blue square and open red triangle) wavelength absorbances of the plasmid samples. Plasmid purification methods included Qiagen MaxiPrep Kit (Non-EF) and Qiagen EndoFree MaxiPrep Kit with one endotoxin removal step (1x EF) or two endotoxin removal steps (2x EF). Additional salt removal using centrifugal filter-based purification was applied to half of the samples (+Filter purification). **(B)** Final dilution factor of the plasmids in the electroporation reaction. **(C)** Percentage of viable T cells pre-electroporation (Pre-EP) and 24 hours post-electroporation among CD8+ and CD8-T cells (n = 3 donors for each, mean ± SD). **(D)** Frequency of CD34+ cells among CD8+ and CD8-T cells at 72 hours post-electroporation (n = 3 donors for each, mean ± SD). **(E)** Mean fluorescence intensity (MFI) of CD34 in CD8+ and CD8-T cells at 72 hours post-electroporation (n = 3 donors for each, mean ± SD). **(F)** Contour plots of CD34 expression at 72 hours post-electroporation among CD8+ (*left*) and CD8-(*right*) T cells for a representative sample in each condition.

**Supplementary Figure 3.**
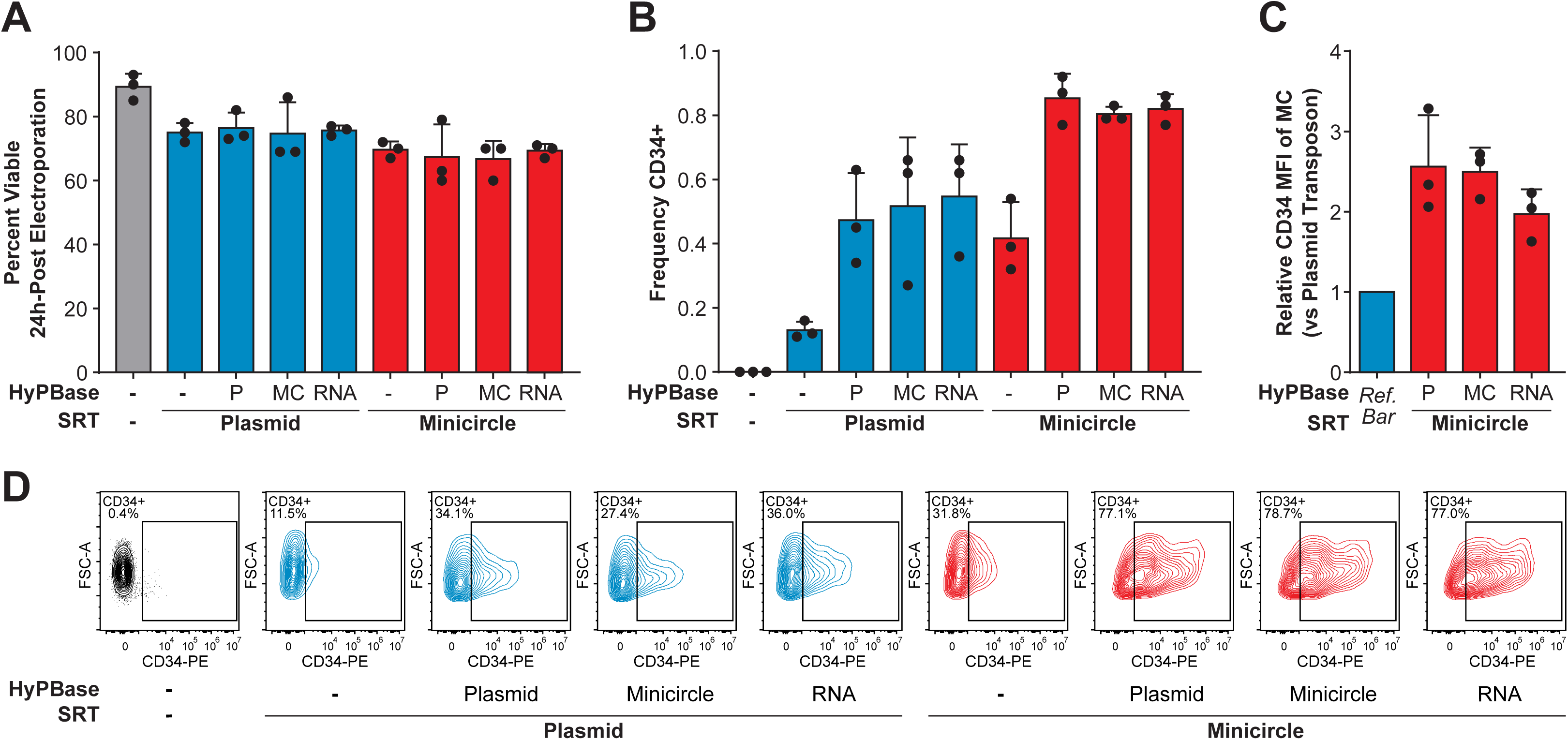
Comparison of HyPBase and SRT-tCD34 forms on transposition efficiency. **(A)** Percentage of viable CD8 T cells at 24 hours post-electroporation of the plasmid (P), minicircle (MC), or RNA forms of HyPBase with the plasmid or minicircle forms of the SRT-tCD34 vector. **(B)** Frequency of CD34+ CD8 T cells at 72 hours post-electroporation of the HyPBase and SRT-tCD34 constructs. **(C)** Relative mean fluorescence intensity (MFI) of CD34 in CD8 T cells at 72 hours post-electroporation with the minicircle SRT-tCD34 vector normalized to the plasmid SRT-tCD34 vector when co-electroporated with the plasmid (P), minicircle (MC), or RNA forms of HyPBase. A bar set at 1 was included as a reference (Ref. Bar). **(D)** Representative contour plots of CD34 expression in CD8 T cells at 72 hours post-electroporation with the various forms of HyPBase and SRT-tCD34 constructs.

**Supplementary Figure 4.**
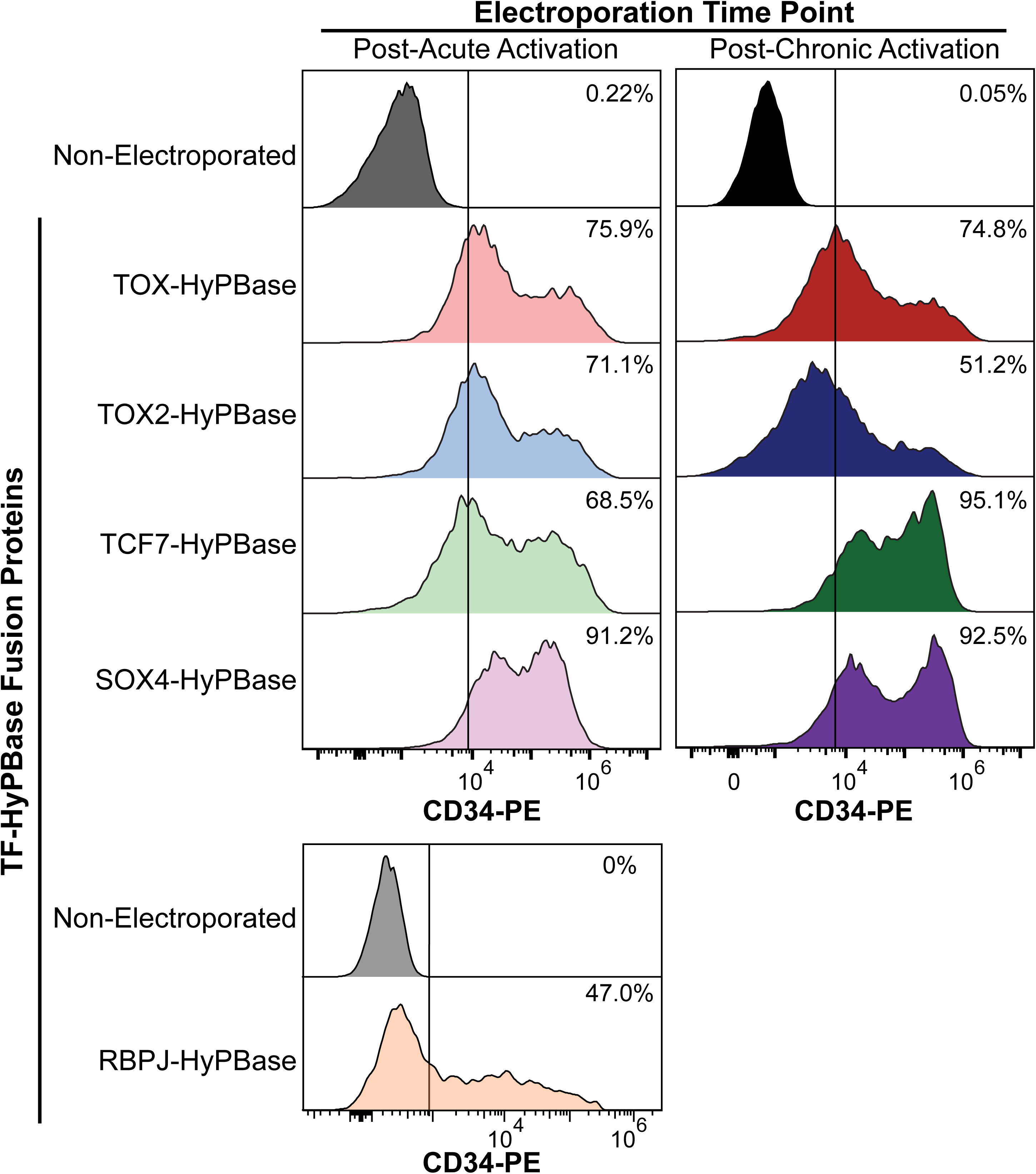
Efficiency of the TF-HyPBase fusion proteins in primary human CD8 T cells. Representative histograms of CD34 expression from the SRT-tCD34 transposon, normalized to the mode, at 72 hours post-electroporation of the TF-HyPBase fusion proteins and SRT-tCD34 vector within *in vitro* acute or chronic activation conditions.

**Supplementary Figure 5.**
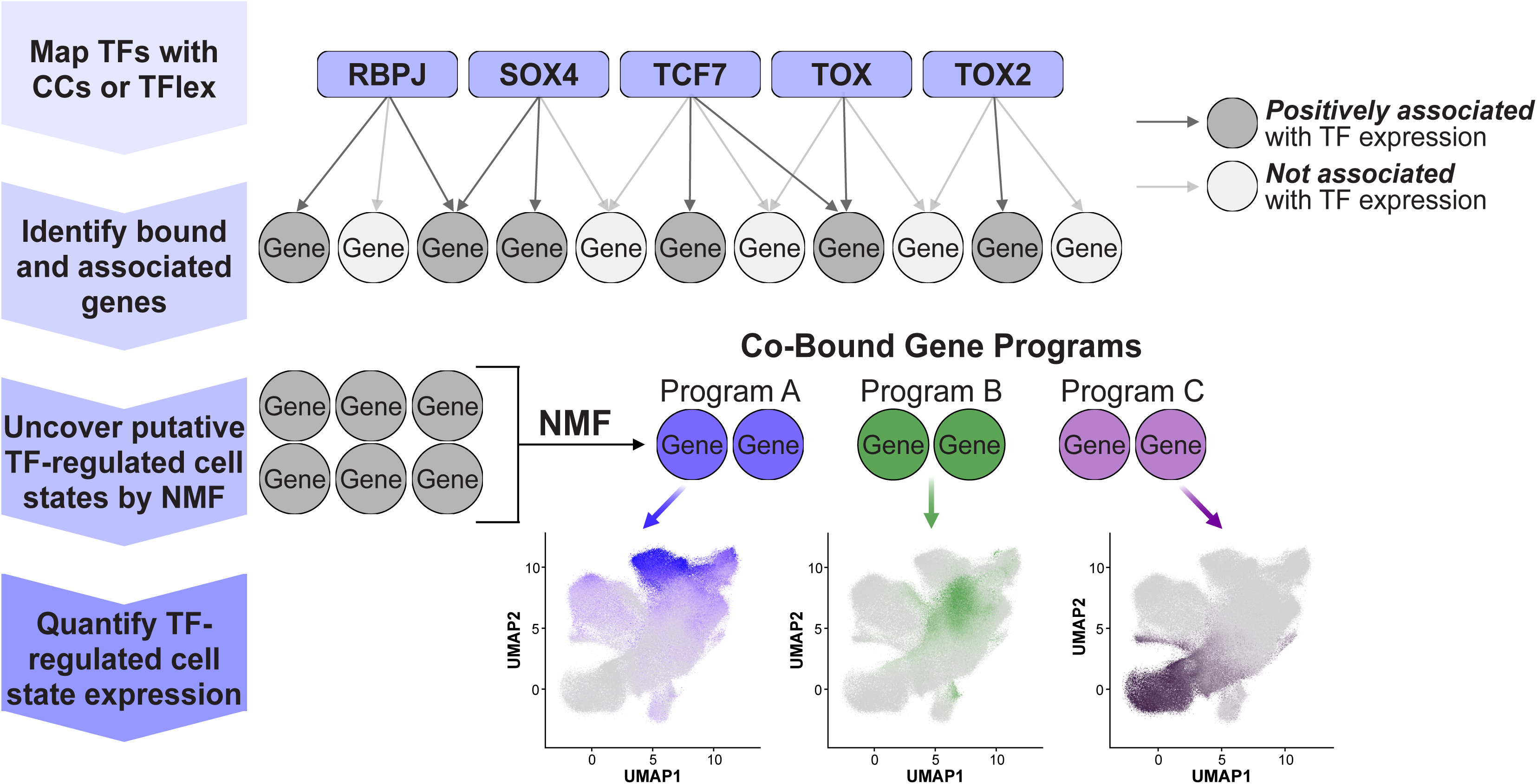
Analytical framework to integrate transcription factor binding data with single cell RNA-sequencing. TF binding sites are first defined by CCs or TFlex. From those binding sites, genes bound at the promoter or an annotated enhancer are identified. Using the scRNA-seq data, the bound genes are subset to those with expression positively associated with the TF to obtain the final bound and associated gene set. All unique genes across the bound and positively associated gene sets are used as input for non-negative matrix factorization (NMF) to uncover co-bound gene programs, representing putative TF-regulated cell states. Lastly, the expression of the TF-regulated cell states is quantified per-gene and as a module score.

**Supplementary Figure 6.**
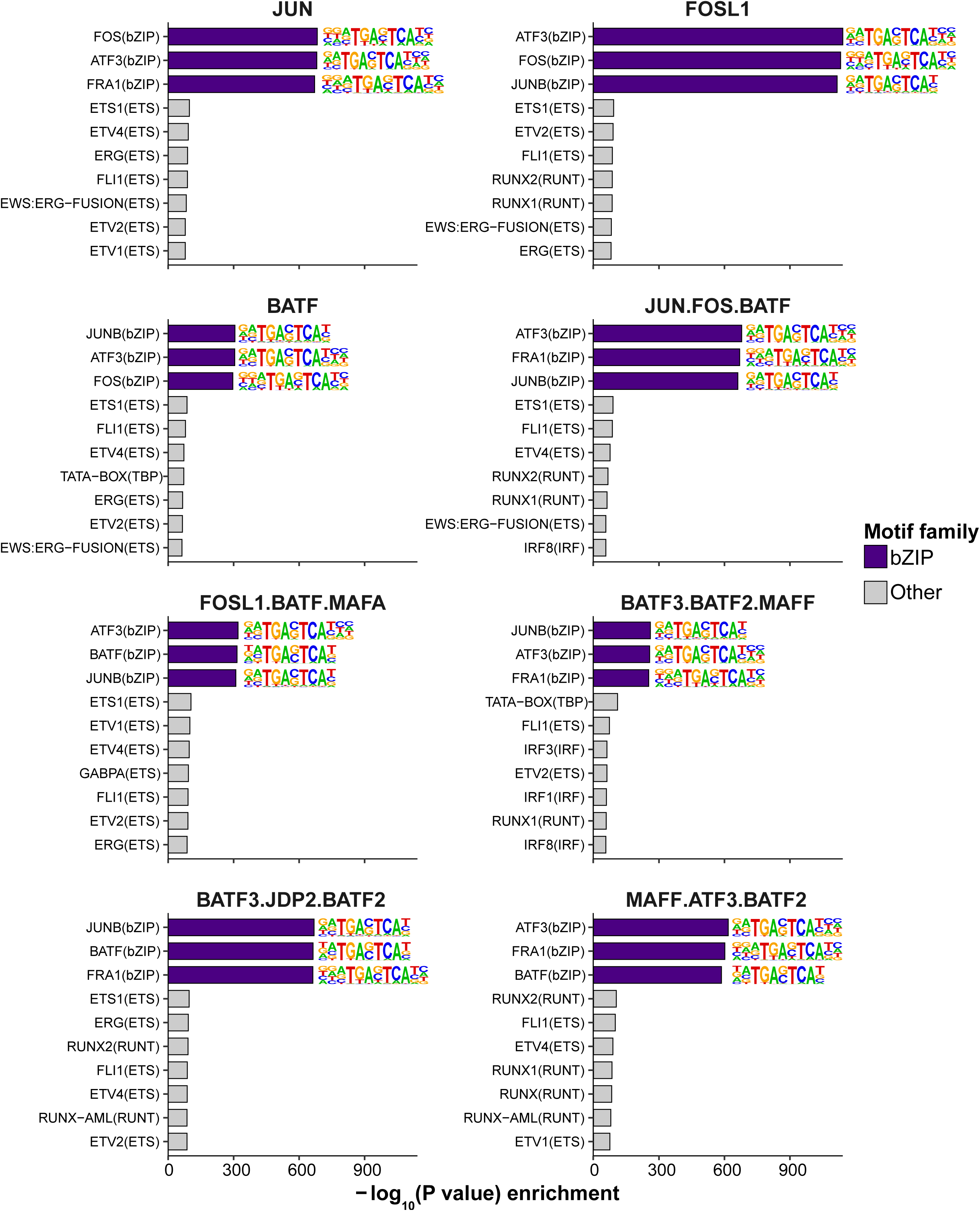
Motif enrichment for the binding sites of natural and domain-swapped bZIP transcription factors defined by TFlex. -log_10_(P value) of HOMER motif enrichment analysis within each TF’s peak set. The enrichment of bZIP motifs (colored dark blue) is expected as all TFs possess a bZIP DNA-binding domain. Only the top three bZIP motifs for each TF are shown because the bZIP motif sequences are highly similar.

## METHODS

### T cell isolation and culture

For the standard CCs data, human adult peripheral blood was obtained from leukocyte reduction system cones that are classified as non-human research under the Washington University Human Research Protection Office. For TFlex samples, human adult peripheral blood was obtained from the Gulf Coast Regional Blood Center (Houston, Texas). Peripheral blood mononuclear cells (PBMCs) were isolated with Ficoll-Paque density gradient medium (Millipore-Sigma, GE17-1440-02) using SepMate-50 tubes (StemCell Technologies, 85460). CD8+ cells were isolated using positive selection with CD8 MicroBeads (Miltenyi Biotech, 130-045-201).

Isolated cells were plated at 1×10^6^ cells mL^−1^ in ImmunoCult-XF T Cell Expansion medium (StemCell Technologies, 10981) supplemented with 1% penicillin-streptomycin (Corning, 30-002-CI). In the acute activation condition, cells were activated for 72 h on plates coated with 5 μg mL^−1^ anti-CD3 (Biolegend, clone OKT3, 317348) and 2 μg mL^−1^ anti-CD28 (Biolegend, clone CD28.2, 302943), and media was supplemented with 300 U mL^−1^ interleukin-2 (IL-2, StemCell Technologies, 78036.1), 10 ng mL^−1^ IL-7 (StemCell Technologies, 78053.1), and 10 ng mL^−1^ IL-15 (StemCell Technologies, 78031.1). In the chronic activation condition, cells were activated with plates coated with 5 μg mL^−1^ anti-CD3 and 2 μg mL^−1^ anti-CD28 for 72 h. Then, cells were re-plated on plates coated with 5 μg mL^−1^ anti-CD3 at 0.5–1.0×10^6^ cells mL^−1^ every 48 h in media supplemented with 5 μM beta-mercaptoethanol (Sigma-Aldrich, M3148) and 300 U mL^−1^ IL-2 (StemCell Technologies, 78036.1) for eight additional days for 11 total days of activation.

### Measurement of population doublings of acutely or chronically activated CD8 T cells

On days 3, 5, 7, 9, and 11 following the initial activation, cells were collected from the wells and resuspended in culture media. Viable cells were counted on the basis of 0.4% trypan blue exclusion using the Countess 3 Automated Cell Counter (Invitrogen, AMQAX2000). Population doublings were calculated by the following equation: ln((Day N Count) / (Day 0 Count)) / ln(2).

### Flow cytometry of acutely or chronically activated CD8 T cells

On day 11 post-initial activation, the cells activated acutely and chronically were cultured on plates coated with 5 μg mL^−1^ anti-CD3 (Biolegend, clone OKT3, 317348) for 48 h in ImmunoCult-XF T Cell Expansion medium (StemCell Technologies, 10981) supplemented with 5 μM beta-mercaptoethanol (Sigma-Aldrich, M3148) and 300 U mL^−1^ IL-2 (StemCell Technologies, 78036.1). A portion of the acutely activated cells remained resting. After 24 h of additional culture, cells were collected, washed in PBS (Sigma-Aldrich, P3813), and stained with Zombie NIR viability dye (1:4,000, Biolegend, 423105) followed by washing in PBS supplemented with 2% fetal bovine serum (FBS, Peak Serum, PS-FB1). Subsequently, cells were stained in PBS supplemented with 2% FBS for 30 min at room temperature with CD39-BV421 (1:100, Biolegend, clone A1, 328214), CD8-BV510 (1:200, Biolegend, clone SK1, 344732), CD3-BV785 (1:200, Biolegend, clone UCHT1, 300472), PD-1-PE/Dazzle594 (1:200, Biolegend, clone EH12.2H7, 329939), TIM-3-PE/Cy7 (1:100, Biolegend, clone F38-2E2, 345013), and CD137-APC (1:100, Biolegend, clone 4B4-1, 309809). Data were acquired on the Cytoflex SRT instrument (Beckman Coulter). Compensation was performed using UltraComp eBeads Compensation Beads (Invitrogen, 01-2222-41) stained with the same concentration of each antibody at 4°C for 30 min. The flow cytometry results were analyzed using FlowJo™ v10.10.0 Software (BD Life Sciences).

### RNA extraction

Total RNA was extracted using the RNeasy Mini Kit (Qiagen, 74104) following the manufacturer’s instructions. Cell pellets were homogenized by vortexing and pipetting in 600 μL buffer RLT supplemented with 1% 2-mercaptoethanol (Sigma-Aldrich, M3148). On-column DNA digest was performed with RNase-free DNase I (Qiagen, 79254). RNA was eluted in nuclease-free water, and the concentration was determined by the Qubit RNA XR assay kit (Invitrogen, Q33224).

### Reverse transcription to generate cDNA from acutely or chronically activated CD8 T cells for quantitative PCR

A reverse transcription reaction was assembled with 1 μg RNA, 2 μL of 50 μM oligo dT primer (Invitrogen, AM5730G), 1 μL of 10 mM UltraPure PCR Deoxynucleotide Mix (Takara, 639125), and nuclease-free water to 14 μL followed by incubating at 65°C for 5 min. After placing on ice for 1 min, 4 μL of 5X reverse transcription buffer (Thermo Scientific, EP0752), 1 μL of RNaseOUT (40 U, Invitrogen, 10777019), 0.5 μL Maxima H minus reverse transcriptase (100 U, Thermo Scientific, EP0752), and 0.5 μL nuclease-free water was added. The reaction was incubated at 50°C for 60 min and 85°C for 10 min. The cDNA was purified with the NucleoSpin Gel and PCR Clean-Up kit (Takara, 740609), and the concentration was determined with the Qubit ssDNA assay kit (Invitrogen, Q10212).

### Quantitative PCR

Quantitative PCR was performed in technical triplicates for each gene. Each reaction was composed of 10 ng cDNA, 0.5 μL of 5 μM forward primer, 0.5 μL of 5 μM reverse primer, 5 μL PowerUp SYBR Green Master Mix for qPCR (Applied Biosystems, A25742), and nuclease-free water to 10 μL. *ACTB* expression was used as the internal control. Reactions were assembled in the Roche-type 384-well full-skirted PCR plate (Midwest Scientific, PR-PCR33842). Quantitative PCR was performed on the QuantStudio 5 Real-Time PCR System (Applied Biosystems, A28570) and analyzed in the QuantStudio Design and Analysis Software (v1.5.3). ROX was used as the passive reference. Quantitative PCR primer sequences are listed in **Supplementary Table 8**.

### Construction of SRT-tCD34 vectors with 25 barcodes for standard CCs

The standard CCs experiments used the minicircle SRT-tCD34 vector with 25 barcodes consisting of 4 bp at the 3’ end of the terminal repeat directly preceding the terminal repeat-genomic DNA junction as previously described^48^. The 25 barcodes were inserted into the plasmid using PCR where the primers encoded the barcodes. Two PCR reactions were assembled. One reaction consisted of 12.5 μL CloneAmp HiFi PCR Premix (Takara, 639298), 0.75 μL of 10 μM Standard_CCs_25_barcode_SRT_pool_Fw_1, 0.75 μL of 10 μM Standard_CCs_25_barcode_SRT_pool_Rv_1, 2 ng P60 Minicircle Standard CCs SRT-tCD34 (plasmid and vector sequences in **Supplementary Table 8**), 1.25 μL dimethyl sulfoxide (DMSO, Sigma-Aldrich, D2650), and nuclease-free water to 25 μL. The second PCR reaction consisted of 12.5 μL CloneAmp HiFi PCR Premix (Takara, 639298), 0.75 μL of 10 μM Standard_CCs_25_barcode_SRT_pool_Fw_2, 0.75 μL of 10 μM Standard_CCs_25_barcode_SRT_pool_Rv_2 (one primer for each of 25 barcodes), 2 ng P60 Minicircle Standard CCs SRT-tCD34 (plasmid and vector sequences in **Supplementary Table 8**), and nuclease-free water to 25 μL. Thermocycling parameters were as follows: 35 cycles of 98°C for 10 s, 55°C for 5 s and 68°C for 25 s. PCR products were incubated with 20 U DpnI (NEB, R0176L) at 37°C for 1 h to digest the input template vector and then purified with the NucleoSpin Gel and PCR Clean-Up kit (Takara, 740609). The PCR products were cloned using the In-Fusion Snap Assembly master mix (Takara, 638948), and the In-Fusion reaction was transformed into Stellar competent cells (Takara, 636763), which were plated onto LB-agar (Lambda Biotech, C121) plates with 50 µg mL^−1^ kanamycin. Full plasmid sequencing was performed using nanopore sequencing (Plasmidsaurus).

### Construction of TFlex SRT-tCD34 vector pools with a distinct sample barcode and diverse SRT barcodes

The sample barcodes and highly diverse SRT barcodes in the TFlex SRT-tCD34 vectors were inserted by PCR. The PCR reaction consisted of 12.5 μL CloneAmp HiFi PCR Premix (Takara, 639298), 0.75 μL of 10 μM TFlex_SRT_Fw, 0.75 μL of 10 μM TFlex_SRT_Rv, 2 ng of P167 minicircle TFlex SRT-tCD34 vector, 1.25 μL DMSO (Sigma-Aldrich, D2650), and nuclease-free water to 25 μL (see **Supplementary Table 8** for vector and primer sequences). Each TFlex_SRT_Rv primer was ordered from Integrated DNA Technologies (IDT) with a single sample barcode and 15 “N” nucleotides to introduce highly diverse SRT barcodes (**Supplementary Table 8**). One reaction was used for each TFlex_SRT_Rv primer, which contain a unique sample barcode and a stretch of 15 random “N” nucleotides which encode the diverse SRT barcodes. Thermocycling parameters were as follows: 35 cycles of 98°C for 10 s, 62°C for 5 s and 68°C for 60 s. PCR products were incubated with 20 U DpnI (NEB, R0176L) to digest the input template plasmid for 1 h and then purified with the NucleoSpin Gel and PCR Clean-Up kit (Takara, 740609). 100 ng of the PCR product was cloned using the In-Fusion Snap Assembly master mix (Takara, 638948) in a 5 μL reaction. Next, 5 μL of the In-Fusion reaction was transformed into 100 μL of Stellar competent cells (Takara, 636763) and plated onto LB-agar (Lambda Biotech, C121) plates with 50 µg mL^−1^ kanamycin. The next day, the colonies were scraped from the LB-agar plate with a cell scraper and cultured in LB-Miller broth (Sigma-Aldrich, L3522) for 14–16 h. Bacterial glycerol stocks were prepared from the overnight culture. Sample barcode identity and diversity in the unique transposon identifier were confirmed with full plasmid sequencing performed using nanopore sequencing (Plasmidsaurus).

### Minicircle SRT-tCD34 vector preparation

SRT-tCD34 was subcloned into the backbone of the pMC.CMV-GFP-SV40PolyA (System Biosciences, MN601A-1) vector in place of the CMV-GFP-SV40PolyA cassette. The minicircle SRT-tCD34 vector was transformed into the minicircle producer strain ZYCY10P3S2T (System Biosciences, MN900A-1). The culture was grown for 13–15 h at 30°C at 250 rpm. Then 1.25 volumes of LB-Miller broth (Sigma-Aldrich, L3522) was added along with a final concentration of 0.02% L-(+)-arabinose (Sigma-Aldrich, A3256) and 40 mM sodium hydroxide (NaOH, Sigma-Aldrich, 221465). The culture was incubated at 37°C at 250 rpm for an additional 5–6 h, after which minicircle DNA was isolated using an endotoxin-free maxi prep (Qiagen, 12362). Resulting DNA was digested for 16 h with NdeI (NEB, R0111L) and RecBCD (NEB, M0345L) in Buffer 4 (NEB, B7004S) supplemented with 1 mM ATP (NEB, P0756S) to remove the parental plasmid and contaminating genomic DNA. The enzymes were heat inactivated by incubating at 70°C for 30 min. The digest reaction was then brought up to 12 mL with a solution of 750 mM sodium chloride (NaCl, Sigma-Aldrich, 746398) and 50 mM 3-(N-Morpholino)propanesulfonic acid (MOPS, Millipore-Sigma, 475922) adjusted to pH 7.0. The endotoxin-free maxi prep (Qiagen, 12362) protocol was repeated at the step of adding 0.1 volume buffer ER. Following isopropanol precipitation, salts were removed by washing the DNA with nuclease free water using a 30 kDa molecular weight cut-off Amicon Ultra-0.5 Centrifugal Filter (Millipore-Sigma, UFC503024). The DNA concentration was quantified with the Qubit 1X dsDNA Broad Range kit (Invitrogen, Q33265). Purity was assessed on the NanoDrop One instrument (Thermo Scientific, ND-ONE-W) based on the 260/230 nm and 260/280 nm wavelength absorbance ratios.

### TF-HyPBase vector preparation

Full information on the source, transcript version, and sequence of the TF-HyPBase fusions is provided in **Supplementary Table 8**. The natural and domain-swapped bZIP TFs were obtained from Ansuman T. Satpathy. The open reading frame of the TFs was cloned into a BsmI restriction site, which was followed by a linker and HyPBase, using the In-Fusion Snap Assembly master mix (Takara, 638948) according to the manufacturer’s protocol. Full plasmid sequencing of the TF-linker-HyPBase vector was performed using nanopore sequencing (Plasmidsaurus).

### *In vitro* transcription of mRNA encoding the TF-HyPBase fusion protein

For the standard CCs experiments, mRNA was prepared using the HiScribe T7 ARCA mRNA kit with PolyA tailing (NEB, E2060S) according to the manufacturer’s protocol. For the TFlex experiments, mRNA was prepared with the HiScribe T7 mRNA Kit with CleanCap Reagent AG kit (NEB, E2080S) according to the manufacturer’s protocol. *E. coli* PolyA polymerase (NEB, M0276S) was used to add a Poly(A) tail to the mRNA. The mRNA was purified with the Monarch spin RNA cleanup kit (NEB, T2040L). mRNA size, quality, and polyA tailing was assessed on the Agilent TapeStation 4200 System (Agilent, G2991BA) using RNA ScreenTape (Agilent, 5067-5576) and RNA sample buffer (Agilent, 5067-5577). Template DNA for *in vitro* transcription was prepared with PCR using the CloneAmp HiFi PCR master mix (Takara, 639298) and plasmid DNA encoding the TF-HyPBase fusion. The T7 promoter sequence was either in the TF-HyPBase fusion protein vector or appended to the 5’ end of the sequence with primers used in PCR to generate the *in vitro* transcription template.

### T cell electroporation

T cells were electroporated using the Neon Transfection System (Invitrogen, MPK5000). T cells were collected on day 3 post-activation for the acute activation condition and day 11 post-continuous activation for the chronic activation condition. Cells were washed in magnesium- and calcium-free PBS (Corning, 21-040-CM) three times and resuspended in buffer R at 40-50×10^6^ cells mL^−1^. The mRNA of unfused HyPBase background control or a TF-HyPBase fusion was added at 25 μg mL^−1^ and the minicircle SRT-tCD34 vector was added at 60 μg mL^−1^. Cells were electroporated in buffer E2 using the 1600 V, 10 ms, and 3 pulse settings. After electroporation, cells were immediately plated in 8 mL pre-warmed ImmunoCult-XF T Cell Expansion medium (StemCell Technologies, 10981) with the same cytokines used for culturing but without penicillin-streptomycin in 10 cm tissue culture dishes. After overnight incubation, viability was assessed using 0.4% trypan blue exclusion and the Countess 3 Automated Cell Counter (Invitrogen, AMQAX2000). Cells were refreshed into ImmunoCult-XF T Cell Expansion medium (StemCell Technologies, 10981) with cytokines and 1% penicillin-streptomycin and plated at a density of 0.25-0.50×10^6^ viable cells mL^−1^. The chronic activation conditions were cultured on plates coated with 5 μg mL-1 anti-CD3 (Biolegend, clone OKT3, 317348) and 2 μg mL-1 anti-CD28 (Biolegend, clone CD28.2, 302943). Cells were collected at day 3-post electroporation in both the acute and chronic activation conditions.

### Cell sorting of SRT-tCD34-positive CD8 T cells

Cells were washed in PBS (Sigma-Aldrich, P3813) and stained with Zombie NIR viability dye (1:4,000, Biolegend, 423105) followed by washing in PBS supplemented with 2% FBS (Peak Serum, PS-FB1). Subsequently, cells were stained with PE anti-human CD34 (1:20, Biolegend, clone 581, 343505) in PBS supplemented with 2% FBS for 30 min at room temperature. Cells were washed once with PBS supplemented with 2% FBS prior to sorting. Cell sorting of viable (Zombie NIR dye-negative), CD34-positive cells was performed on the CytoFLEX SRT instrument (Beckman Coulter). The flow cytometry results were analyzed using FlowJo™ v10.10.0 Software (BD Life Sciences).

### Reverse transcription of RNA to generate cDNA for standard CCs and TFlex libraries

A reverse transcription reaction was assembled with 5 μg RNA, 0.5 μL of 100 μM SMART_dT18VN primer for standard CCs or SMART_Nest_dT18VN for TFlex, 1 μL of 10 mM UltraPure PCR Deoxynucleotide Mix (Takara, 639125), and nuclease-free water to 14 μL. The reaction was incubated at 65°C for 5 min and then placed on ice for 1 min. Then, 4 μL of 5X reverse transcription buffer (Thermo Scientific, EP0752), 1 μL of RNaseOUT (40 U, Invitrogen, 10777019) for standard CCs or 1 μL SUPERaseIn RNase Inhibitor (1 U, Invitrogen, AM2694) for TFlex, and 1 μL Maxima H minus reverse transcriptase (200 U, Thermo Scientific, EP0752) was added. The reaction was incubated at 50°C for 60 min for standard CCs or 45 min for TFlex followed by 85°C for 10 min. 1 μL RNase H (5 U, NEB, M0297S) was then added and incubated for 37°C for 30 min to degrade RNA-DNA hybrid molecules. For TFlex, cDNA was cleaned-up with the NucleoSpin Gel and PCR Clean-Up kit (Takara, 740609) according to the manufacturer’s protocol to remove primers and RNA. The cDNA concentration was determined with the Qubit ssDNA assay kit (Invitrogen, Q10212). Primer sequences are listed in **Supplementary Table 8**.

### Amplification of SRT-derived transcripts for standard CCs and TFlex libraries

Standard CCs used a single round of PCR to amplify SRT-derived transcripts. A reaction of the following components was assembled: 100 ng of cDNA, 12.5 μL KAPA HiFi HotStart ReadyMix PCR kit (Roche Diagnostics, 07958927001), 0.5 μL of 25 μM SMART primer, 0.5 μL of 25 μM Standard_CCs_SRT_Amp primer, and nuclease-free water to 25 μL. Thermocycling parameters were as follows: 95°C for 3 min; 20 cycles of 98°C for 20 s, 65°C for 30 s, 72°C for 5 min; 72°C for 10 min. Amplified transposon-derived transcripts were purified with a 0.6X AMPure XP beads clean-up (Beckman Coulter, A63880). The concentration of the purified PCR products was obtained with the Qubit 1X dsDNA High Sensitivity assay kit (Invitrogen, Q33231).

TFlex used two rounds of nested PCR to amplify SRT-derived transcripts. First, 100 ng of cDNA was input into a PCR reaction with 12.5 μL KAPA HiFi HotStart ReadyMix PCR kit (Roche Diagnostics, 07958927001), 0.75 μL of 10 μM SMART primer, 0.75 μL of 10 μM V3_SRT_Amp primer, and nuclease-free water to 25 μL. Thermocycling parameters were as follows: 95°C for 3 min; 12 cycles of 98°C for 20 s, 65°C for 15 s, 72°C for 6 min; 72°C for 6 min. Amplified transposon-derived transcripts were purified with a 0.5X SPRI bead clean-up (Beckman Coulter, B23317). DNA was eluted in 14 μL of nuclease-free water, and 11 μL was input into a second PCR reaction with 12.5 μL KAPA HiFi HotStart ReadyMix PCR kit (Roche Diagnostics, 07958927001), 0.75 μL of 10 μM SMART_Amp_2 primer, 0.75 μL of 10 μM V2_LacZ_SRT_Amp primer. Thermocycling parameters were as follows: 95°C for 3 min; *10 or 12* cycles of 98°C for 20 s, 65°C for 15 s, 72°C for 6 min; 72°C for 6 min. The concentration of the purified PCR products was obtained with the Qubit 1X dsDNA High Sensitivity assay kit (Invitrogen, Q33231). Libraries were prepared from both the 10 and 12 cycle reactions, and the reads from each were combined. Primer sequences are listed in **Supplementary Table 8**.

### Final library preparation and sequencing for standard CCs and TFlex libraries

Standard CCs libraries were prepared using the Nextera XT DNA Library Preparation Kit (Illumina, FC-131-1024). 1 ng of the amplified transposon-derived transcripts was combined with 10 μL Tagment DNA (TD) Buffer, 5 μL Amplicon Tagment Mix (ATM) Buffer, and nuclease-free water to 15 μL. The reaction was incubated at 55°C for 8 min followed by a 10°C hold. The tagmentation reaction was stopped by the addition of 5 μL Neutralization Tagment (NT) Buffer and incubation at room temperature for 5 min. Dual-index sequences were appended in a PCR reaction consisting of 15 μL Nextera PCR Mix (NPM), 8 μL nuclease-free water, 1 μL of 10 μM a custom Standard_CCs_i5_library_amplification, and 1 μL of 10 μM of the standard Nextera N7 indexed primer. The following thermocycling parameters were used: 72°C for 3 min; 95°C for 30 s; 15 cycles of 95°C for 10 s, 55°C for 30 s, 72°C for 30 s; 72°C for 5 min. Following amplification, PCR products were purified with a 0.7X AMPure XP beads (Beckman Coulter, A63880) purification. The libraries were eluted in 11 μL nuclease-free water and quantified on the Agilent TapeStation 4200 System (Agilent, G2991BA) using the High Sensitivity D5000 ScreenTape (Agilent, 5067-5592) and reagents (Agilent, 5067-5593).

TFlex libraries were also prepared using the Nextera XT DNA Library Preparation Kit (Illumina, FC-131-1024). However, the following modifications were made to the standard CCs library: tagmentation was done for 5 min instead of 8 min at 55°C, library amplification used the custom TFlex_i5_library_amplification primer in place of the Standard_CCs_i5_library_amplification primer, amplification used 12 cycles instead of 15 cycles; and two consecutive 0.56X SPRI bead (Beckman Coulter, B23317) clean-ups were performed following library amplification. Primer sequences are listed in **Supplementary Table 8**.

The standard CCs and TFlex libraries were quantified on the Agilent TapeStation 4200 System (Agilent, G2991BA) using the High Sensitivity D5000 ScreenTape (Agilent, 5067-5592) and reagents (Agilent, 5067-5593). Sequencing was performed at the Genome Access Technology Center at the McDonnell Genome Institute (4444 Forest Park Ave., St. Louis, MO 63108) on the NovaSeq 6000 or NovaSeq X Plus sequencing instrument with 2×150 bp reads.

### Standard CCs sequencing data pre-processing

FASTQ paired-end reads were filtered for R1 reads with the structure 5’-[terminal repeat sequence] [4 bp SRT barcode] [HyPBase insertion site “TTAA”] [genomic DNA]. Then, cutadapt (v2.10) was used to filter and trim reads containing the 5’ terminal repeat sequence “CGTCAATTTTACGCAGACTATCTTT” with an exact match and no indels. Next, the umi-tools (v1.0.0) *extract* function was used to capture and trim the SRT barcode, matching a whitelist of supplied 4 bp SRT barcodes. Finally, cutadapt (v2.10) was applied to select and trim reads containing “GTTAA,” where “G” is last base pair of the terminal repeat and “TTAA” is the HyPBase insertion site motif, as well as trim any trailing adapter sequences. Reads were aligned to human reference genome hg38 using bowtie2 (v2.4.2) with default parameters and filtered for mapped primary alignments. Custom python scripts using pysam were used to output unique insertion sites, read count, strand, and SRT barcode in qBED format^71^. Each sample of standard CCs received approximately 10-15 million final processed reads.

### TFlex sequencing data pre-processing

FASTQ paired-end reads were filtered for pair-end reads with R1 structure of 5’-[PCR adapter sequence] [10 bp sample barcode] [15 bp SRT barcode] and R2 structure of 5’-[genomic DNA] [HyPBase insertion site “TTAA”] [terminal repeat]. cutadapt (v2.10) was used to trim paired reads where the R1 containing the 5’ PCR adapter sequence (-g “TAGCTATCCTTCGCAAGACCCTTC”) with up to 2 mismatch and no indels, while similarly R2 read were trimmed up at the TTAA-terminal repeat junction (-A “TTAACCCTAG”) using an exact match with minimum length of 10 bp. However, untrimmed R2 reads that did not hit the terminal repeat (genomic DNA only) were retained. Next, the umi-tools (v1.0.0) *extract* function was then used to capture the 10 bp sample barcode and 15 bp SRT barcode from the R1 read. R2 reads were then aligned to human reference genome hg38 using bowtie2 (v2.4.2) with default parameters and filtered for mapped primary alignments. Custom python scripts using pysam were used to output unique insertion site, read count, strand, and “library/SRT sample barcode/SRT barcode” in qBED format^71^. Each sample barcode within the multiplexed TFlex libraries received approximately 10-15 million final processed reads.

### Demultiplexing of pooled samples, SRT barcode deduplication, and SRT insertion site assignments in TFlex

The output qBED file contained one row for each fragment, which is each unique combination of the sample barcode and SRT barcode from R1 and the end position of the R2. The R2 sequence is oriented such that the read is moving towards the junction of the SRT and genomic DNA, which represents the TF binding site. However, not all R2 reads reach junction of the SRT and genomic DNA due to variation in the size of the fragments in the final library. Additionally, each SRT transcript-derived molecule from the SRT insertions will be associated with multiple fragments arising from fragmentation during the library preparation. We therefore devised a strategy to use R2 to assign the genomic position of SRT insertions and deduplicate multiple fragments associated with the same SRT insertion.

First, sequencing errors in the sample barcodes were corrected against a whitelist with a Hamming distance threshold of two. Samples were then demultiplexed based on the sample barcode. Next, for each sample, fragments within the same genomic region that possessed the same sample and SRT barcodes were deduplicated. This was performed by first using the CCaller method of pycallingcards (v1.0.0)^72^ to group the fragments into variable width genomic bins such that all fragments of the same sample and SRT barcodes were within the same genomic bin. Bins with fewer than five fragments were discarded to reduce noise in the data. Within each bin, SRT barcode sequencing errors were corrected using the directional method from UMI-tools (v1.1.6)^3^ with a Hamming distance threshold of one. Next, after correction, fragments of the same sample and SRT barcodes within each bin were assigned to the coordinate of the fragment that was most proximal to the junction of the SRT and genomic DNA. The coordinate closest to the SRT-genomic DNA junction was the most upstream coordinate for positive strand SRT insertions or the most downstream coordinate for SRT insertions on the negative strand. After position assignment, fragments with the same sample barcode, SRT barcode, and genomic coordinate were collapsed into one to deduplicate them, yielding the final position and unique SRT insertion counts per genomic coordinate. Lastly, the final deduplicated SRT insertion data for all samples of the same TF were combined to generate the final processed qBED file.

### Peak calling for standard CCs and TFlex libraries

The final processed qBED files for each TF and the unfused HyPBase background control for standard CCs or TFlex were input into SPAN2.0^73^ (v2.0.6652), a semi-supervised peak calling method, using a bin size of 50 bp, FDR cutoff of 0.01, and no background control. Resulting peaks with a width less than 200 bp were extended equally on both sides to reach 200 bp, and after extension, peaks within 250 bp of each other were merged into one peak range. These peaks served as preliminary TF binding sites. The final binding sites were determined by first quantifying the SRT insertion counts for the unfused HyPBase background control and all other TFs within each TF’s peak set. The acute activation standard CCs, chronic activation standard CCs, and TFlex data were processed as separate groups. This resulted in an SRT insertion count matrix for each TF of the group where peak ranges were rows and samples were columns. SRT insertion counts for each TF were then normalized by multiplying the counts by a size factor equal to the TF’s total SRT insertion counts divided by the geometric mean of all TFs’ total SRT insertion counts. The normalized count matrix for each TF was used in DESeq2 (v1.46.0)^74^ to determine peaks enriched above the unfused HyPBase background control and differential peaks relative to other TFs, using a local fit to estimate dispersion. The final peak set for each TF enriched above the unfused HyPBase control or other TFs was defined by two cutoffs: log_2_ fold change≥1 of normalized SRT insertions relative to control (unfused HyPBase or another TF) and normalized SRT insertion counts of the TF≥10 for standard CCs or ≥8 for TFlex. Peaks unique to a TF among the dataset were defined as the peaks meeting both filter criteria relative to each other TF and the unfused HyPBase background control.

### Data analysis and visualizations

All data analyses and plots were generated in R (v4.4.1)^75^. All plots were created with the ggplot2 package (v3.5.2)^76^ or GraphPad Prism (v10.6.0), except for heatmaps, which were generated using the ComplexHeatmap package (v2.22.0)^77^. Color palettes were obtained from ArchR (v1.0.2)^78^, RColorBrewer (v1.1-3)^79^, or viridis (v0.6.5)^80^ packages.

### Genome browser visualization of data

All genome browser visualizations were generated using the JBR genome browser (v2.0.6643)^73^ using bigwig files of normalized SRT insertion counts binned in 50 bp windows. SRT insertion counts were normalized by a size factor equal to each TF’s total SRT insertion counts divided by the geometric mean of all TFs’ total SRT insertion counts. Bulk ATAC-seq data were visualized as the average of all samples per cell subset using the bigwigAverage function from deepTools (v3.3.0)^81^. Single cell ATAC-seq data in bigwig format for genome browser visualization was obtained from GSE280508^13^.

### HOMER motif analysis

Motif analysis was performed with HOMER (v5.1)^82^ on the final set of CCs peaks using ‘findMotifsGenome’ (<bed_file_of_CC_peaks> hg38 <output_dir> -size given -mask).

### HOMER peak annotations

The peaks in **Supplementary Table 1** were annotated with the ‘annotatePeaks’ function of HOMER (v5.1)^82^. The peaks meeting the following criteria were input into this function: ≥1 log_2_ fold change of normalized SRT insertions relative to the unfused HyPBase background control and ≥10 normalized SRT insertion counts for standard CCs or ≥ 8 for TFlex.

### Permutation testing of TF binding site enrichment in regions of interest

Permutation-based overlap analyses were performed using regioneR (v1.38.0)^83^. To test for enrichment of TF peaks within ±5 kb transcription start site windows, 1,000 random permutations of TF peak locations across the genome were used to create a null distribution and perform a one-sided test for enrichment. The Hsapiens.UCSC.hg38.knownGene (v3.20.0)^84^ gene annotations were used to define transcription start sites. Statistically significant enrichment of TF binding sites at specified enhancer regions was tested using all bulk ATAC-seq peaks as the background universe and the midpoint of the TF peaks as the query set for overlap with the enhancer regions. The null distribution of overlaps was created with 1,000 permutations of the TF peaks across the universe of the bulk ATAC-seq peaks, and statistical significance was determined with a one-sided test for enrichment.

### Gene ontology (GO) analysis

The nearest protein-coding genes bound ±1 kb of the transcription start site as per the ‘annotatePeaks’ function of HOMER were identified for each TF. These genes were queried for term enrichment in the biological processes categories of GO terms using the clusterProfiler package (v4.14.3)^85^. The top three enriched terms sorted low to high by FDR for each TF were plotted.

### Bulk ATAC-seq data processing of peripheral human CD8 T cell subsets for visualizations

The raw ATAC-seq data peripheral human CD8 T cell subsets from GSE179613^18^ from Gene Expression Omnibus was aligned to the hg38 human reference genome with bowtie2 (v2.3.5.1)^86^ following adaptor trimming with cutadapt (v1.18)^87^. Unmapped, unpaired, and mitochondrial reads were removed using samtools (v1.9)^88^. ENCODE blacklist regions^89^ were removed using bedtools (v2.31.1)^90^ and PCR duplicates were removed using picard (v2.8.1)^91^. Bigwigs normalized for library size were generated for each sample and merged by cell type across all samples within the dataset using the deepTools (v3.3.0)^81^ bigwigAverage function.

### Public bulk RNA-seq and ATAC-seq data of human CD8 T cell processing for analysis

Log_2_-normalized and batch-corrected counts of bulk RNA-seq and ATAC-seq of human CD8 T cells were obtained from Gene Expression Omnibus (GSE179613)^18^. ATAC-seq peak coordinates were lifted over from the hg19 genome to the hg38 genome using the hg19ToHg38.over.chain file (UCSC) with the liftOver function of the rtracklayer package (v1.66.0)^92^. Only the ATAC-seq peaks that mapped to a single hg38 coordinate were retained. The RNA-seq dataset was filtered to retain only the following subsets: CD8_Naive, CD8_EMRA, CD8_EM2, CD8_EM1, CD8_CM, CD8_SCM-R3+, CD8_SCM-R3-, CD8_PD1+CD39+. Lastly, the bulk ATAC-seq and RNA-seq datasets were both filtered to retain only samples that contained both ATAC-seq and RNA-seq data.

### Public CD8 T cell scRNA-seq data processing for analysis

The scRNA-seq atlas raw gene counts matrix and metadata were downloaded from Zenodo (10.5281/zenodo.13382785)^42^. The cells were subset by tissue (peripheral blood mononuclear cells, tumor-infiltrating lymphocytes, and lymph node), disease type (healthy, solid tumor), and percent mitochondrial genes (<7.5%). Cells with the “ag_spcific” value for the subset column were excluded because these cells were from adoptive cell therapy. Otherwise, all data and metadata, including UMAP coordinates and cluster assignments and names, were used as is without further processing.

### Annotation of genes bound at the promoter or enhancer by the TFs

First, to identify candidate enhancer region ATAC-seq peaks, the peak ranges were intersected with the coordinates of the ENCODE-rE2G enhancer region database (processed coordinates in **Supplementary Table 2**)^26^. The genes assigned to the enhancer region ATAC-seq peaks were also obtained from the ENCODE-rE2G enhancer region database^26^. ATAC-seq peaks within a ±5 kb window of a transcription start site, defined by the “promoters” function of GenomicFeatures (v1.58.0)^93^ with TxDb.Hsapiens.UCSC.hg38.knownGene (v3.20.0)^84^ gene annotations, were also annotated as candidate enhancers of the transcription start site-proximal genes. Next, the enhancer region ATAC-seq peak and gene pairs were subset to only those that had an ATAC-seq peak overlapping with a TF binding site in the CCs TF binding data. Lastly, the association between each enhancer region ATAC-seq peak signal with the expression of its corresponding gene was quantified by simple linear regression (*Gene Expression* ∼ *β*1(*ATAC Peak Signal*) + *β*0) using the lm function of the stats (v4.4.1) package and by Spearman correlation using the cor.test function of the stats (v4.4.1) package. The P values were adjusted using Benjamini– Hochberg correction through the p.adjust function of the stats package (v4.4.1)^94^. Only enhancer region ATAC-seq peak-gene pairs meeting these criteria were used for downstream analyses: FDR of linear regression<0.05, *β*1>0.15, R^2^>0.15, and FDR of Spearman correlation<0.05. This final set of co-regulated enhancer-gene pairs was used to annotate enhancer-bound genes for each TF. Lastly, genes bound at the promoter by each TF regardless of overlap with an enhancer region or ATAC-seq peak were defined as genes bound within a ±1 kb window around the transcription start site of genes. All range intersections to find overlaps were performed using the findOverlaps function of GenomicRanges (v1.58.0)^93^. The TF binding annotations for enhancer region ATAC peaks with the association metrics and non-enhancer region promoter-bound genes are in **Supplementary Table 3**. Of note, P values were adjusted using only the subset of enhancer region ATAC-seq peaks bound by TOX, TOX2, TCF7, SOX4, or RBPJ for the main figures or the subset bound by any of the 13 TFs for **Supplementary Table 3**.

### Identification of TF-bound and associated genes

The association between TF expression and each of its promoter- or enhancer-bound genes was determined within the scRNA-seq atlas using metacells to reduce sparsity and a proportionality metric. Metacells were created through the MetacellsByGroups function of the hdWGCNA package (v0.4.06)^95^ by summing the counts of the 15 nearest neighbor single cells per cluster using the UMAP reduction. The metacell count matrix was then used to calculate the proportionality metric of association, which is superior relative to other measures of association for scRNA-seq data as previously described^96,97^. To calculate proportionality, the raw count matrix of the metacells was normalized by a centered log ratio using the NormalizeCounts function of Seurat (v5.2.0)^98^. Next, as previously reported^96^, the following equation was used to calculate the proportionality (*ρ_p_*) between TF expression (*TF*_i_) and the expression of each bound gene (*BoundGene*_j_):

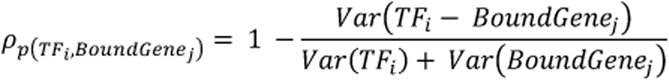

The proportionality was calculated separately in the memory and exhausted subsets as these are distinct phenotypes where the relation of TF expression to its bound gene may be different. The TF-bound genes were then subset to those with |*ρ_p_*| > 0.15 in the memory or exhausted subsets. Genes with opposite *ρ_p_* signs (positive or negative) in the memory and exhausted subsets were removed. Lastly, the final TF bound and associated gene sets were created for both the positively and negatively associated bound genes that met the *ρ_p_* threshold in either the memory or exhausted subsets (**Supplementary Table 5**). The module score for the final bound and positively associated gene sets was quantified using the log-normalized counts of the metacells, which were normalized by the NormalizeCounts function of Seurat (v5.2.0)^98^, and the AddModuleScore_UCell function of UCell (v2.10.1)^99^.

### Identification of NMF-derived TF co-bound programs

Co-bound gene programs were identified by inputting all unique genes across the final TF bound and positively associated gene sets into the runNMF function of the GeneNMF package (v0.9.1)^100^, which uses the RcppML NMF framework (v0.3.7)^101^. First, the log-normalized expression matrix of the metacells from the scRNA-seq atlas was scaled and centered. The matrix was then subset to only the TF bound and positively associated genes. This filtered matrix was input into the runNMF function with k=9 and an L1 (Lasso) regularization of 0.35. The k=9 parameter was chosen because lower k values conflated distinct programs while higher k values led to programs without clear biological relevance. The gene loadings for each program were then normalized to sum to 1, and the genes composing the top 80% of the cumulative normalized NMF loading weights were selected as the final representative gene set for each program. The gene programs were named based on classic, characteristic marker genes. Expression of the final gene programs in the log-normalized matrix of the scRNA-seq atlas metacells was quantified as a module score using the AddModuleScore_UCell function of UCell (v2.10.1)^99^ and on a per-gene level as the mean expression per gene in the metacells of each cluster.

### Standard CCs and TFlex comparison

The final processed SRT insertion counts were summed within 200 bp width genomic bins. The genomic region at chromosome 2:32916200-32916800 was excluded from this analysis in the TFlex samples due to an unusually high SRT insertion count, approximately 10- to 20-fold greater than the next highest insertion count in any other bin. The signal-to-noise ratio was calculated as the ratio of the mean SRT insertion counts in the top 20^th^ percentile of genomic bins ranked by SRT insertion counts to that of the bottom 20^th^ percentile of genomic bins.

### Comparison of JUN, FOSL1, BATF, and JUN.FOS.BATF

All defined peaks of JUN, FOSL1, BATF, and JUN.FOS.BATF were merged using the reduce function of the GenomicRanges package (v1.58.0)^93^ to facilitate visualizations. Next, a matrix of log_2_-transformed normalized SRT insertion counts for each TF in the merged peaks was created and Z-scored across the TFs. The peaks were then manually classified based on which TFs had a positive Z-score value in the peak. For example, the JUN and JUN.FOS.BATF shared peak set was the subset of merged peaks with a positive Z-score value for JUN and JUN.FOS.BATF and a negative Z-score value for FOSL1 and BATF.

Co-bound genes among these TFs were identified using the CCs TF binding and pseudobulked scRNA-seq data for TF overexpression in human CAR T cells (GSE280509)^13^. We identified co-bound genes by first intersecting the JUN.FOS.BATF peaks with the JUN or BATF peak sets using the findOverlaps function of GenomicRanges (v1.58.0)^93^. The overlapping peaks were then associated with genes based on binding at an annotated enhancer region from the ENCODE-rE2G enhancer database^26^ (**Supplementary Table 2**) or at a gene’s promoter (±500 bp of a transcription start site) using the gene annotations from TxDb.Hsapiens.UCSC.hg38.knownGene (v3.20.0)^84^. Lastly, differentially expressed genes were defined in the pseudobulked scRNA-seq data as genes with FDR<0.05 and an absolute log_2_ fold change≥0.75 relative to the control according to DESeq2^74^.

